# SARS-CoV-2 Spike Protein Interacts with Multiple Innate Immune Receptors

**DOI:** 10.1101/2020.07.29.227462

**Authors:** Chao Gao, Junwei Zeng, Nan Jia, Kathrin Stavenhagen, Yasuyuki Matsumoto, Hua Zhang, Jiang Li, Adam J. Hume, Elke Mühlberger, Irma van Die, Julian Kwan, Kelan Tantisira, Andrew Emili, Richard D. Cummings

## Abstract

The spike (S) glycoprotein in the envelope of SARS-CoV-2 is densely glycosylated but the functions of its glycosylation are unknown. Here we demonstrate that S is recognized in a glycan-dependent manner by multiple innate immune receptors including the mannose receptor MR/CD206, DC-SIGN/CD209, L-SIGN/CD209L, and MGL/CLEC10A/CD301. Single-cell RNA sequencing analyses indicate that such receptors are highly expressed in innate immune cells in tissues susceptible to SARS-CoV-2 infection. Binding of the above receptors to S is characterized by affinities in the picomolar range and consistent with S glycosylation analysis demonstrating a variety of N- and O-glycans as receptor ligands. These results indicate multiple routes for SARS-CoV-2 to interact with human cells and suggest alternative strategies for therapeutic intervention.

## Introduction

SARS-CoV-2 is a positive-sense RNA enveloped virus characterized by a surface Spike (S) glycoprotein^1^. During host cell invasion, S binds to receptors on cell membranes, such as angiotensin-converting enzyme 2 (ACE2)^1, 2, 3^. However, the nature and function of the S glycosylation is not fully understood.

Densely glycosylated with multiple Asn-linked (N-glycans) and a few Ser/Thr-linked (O-glycans)^4, 5^, the S protein of SARS-CoV-2 potentially presents ligands for a variety of innate immune receptors, including C-type lectin receptors (CLRs), that are known to bind specific glycans mostly in a Ca^2+^-dependent manner^6, 7^. CLRs such as DC-SIGN/CD209, L-SIGN/CD209L/CLEC4M, mannose receptor/MR/MRC1/CD206, MGL/CLEC10A/CD301, and Dectin-2/CLEC6A, are highly expressed within the human immune system including monocytes, dendritic cells, and macrophages, functioning as the first-line of defense against invading viruses and pathogens^8, 9^. As known pattern-recognition receptors, CLRs, especially DC-SIGN, can direct host immune responses against numerous pathogens in a glycan-specific manner by modulating Toll-like receptor-induced activation ^10^.

Evidence implicates innate immune cells in the pathogenesis of SARS-CoV-2. Over 80% of patients with SARS-CoV-2 infection present with lymphopenia and an increased neutrophil-lymphocyte ratio^11^. Patients with severe COVID-19 exhibit hyperactive macrophages in the bronchoalveolar lavage fluid (BALF) and oropharyngeal swab^12^. Likewise, increased infiltration and activation of macrophages is observed in biopsy or autopsy specimens from COVID-19 patients^13^. Previous studies of the closely-related SARS-CoV demonstrated that primary human monocytes and dendritic cells can be infected^14, 15^, and SARS-CoV S can bind DC-SIGN and L-SIGN^16, 17, 18, 19^. Eight glycosylation sites of SARS-CoV S were identified to be involved in their interactions^20, 21^, among which six are conserved in SARS-CoV-2 (**Supplementary Fig. 1**). However, it is not known whether SARS-CoV-2 interacts with a variety of CLRs.

Here, we demonstrate that many different CLRs directly bind in a glycan-dependent manner to the S glycoprotein of SARS-CoV-2 with picomolar affinities. Binding of DC-SIGN can trigger the internalization of S in 3T3-DC-SIGN+ cells, which implies the potential involvement in viral entry. Furthermore, we dissect the N- and O-glycan sequences on the recombinant SARS-CoV-2 S, and identify glycan features that are crucial for interactions with these CLRs. The analyses of open accessible single-cell RNA sequencing data confirm that various human tissues and their resident immune cells differentially express CLRs, including MR, MGL and DC-SIGN in bronchoalveolar macrophages in patients with SARS-CoV-2. This is in direct contrast to the absence of ACE2 expression within the same cell types across the tissues. Our study identifies new SARS-CoV-2 binding receptors expressed on innate immune cells, particularly on macrophages and dendritic cells, which could accentuate severe pathological inflammation along with cytokine release syndrome. The results suggest potential additional routes for viral infection and new anti-viral strategies.

## Results

### Multiple CLRs bind SARS-CoV-2 S in a glycan-dependent manner

Multiple CLRs including DC-SIGN, L-SIGN, MR (C-type lectin domains 4-7) and MGL exhibited strong binding to the recombinant full-length S produced in human embryonic kidney HEK293 cells (**Fig. 1a-c & e**). HEK293 cells are known to present a spectrum of human glycosylation reflective of the kidney and other epithelial tissues^22^. DC-SIGN, L-SIGN and MGL also bound to recombinant S1, the subunit involved in ACE2 recognition. By contrast, another CLR, Dectin-2, did not bind S or S1, but bound the positive control, a yeast extract (EBY-100) containing mannan-type ligands (**Fig. 1d**). Binding of DC-SIGN, L-SIGN, MR and MGL was glycan-dependent, as it was sensitive to treatment with sequence-specific glycosidases (**Fig. 1f**). Binding of DC-SIGN and L-SIGN was attenuated by Endoglycosidase H (Endo H, oligomannose and hybrid type N-glycan-targeting) and eliminated by PNGase F (N-glycan-targeting), suggesting that the two CLRs bind to S via both oligomannose and complex N-glycans. MR binding was abolished by both Endo H and PNGase F (**Fig. 1c**), suggesting that oligomannose N-glycans are the ligands. A profound reduction in MGL binding was observed by exposure to PNGase F, indicating that MGL ligands reside primarily on N-glycans. Sialic acid appeared to mask some of the MGL ligands as neuraminidase (Neu) treatment slightly increased the binding (**Fig. 1e**), which was similar to the effect on bovine submaxillary mucin (BSM) with abundant O-glycans comprised of sialylated N-acetylgalactosamine (STn antigen). The effects of glycosidase treatment were confirmed by loss or gain of binding by glycan-binding lectins GNA, ConA and VVA (**Supplementary Fig. 2a**, **b & d)**. Interestingly, neither DC-SIGN nor L-SIGN bound recombinant S2 (**Supplementary Fig. 2e-h**), the subunit mediating cell fusion. In all cases CLR binding to S and S1 was Ca^2+^-dependent as expected (**Supplementary Fig. 2i-l**).

**Fig. 1.**
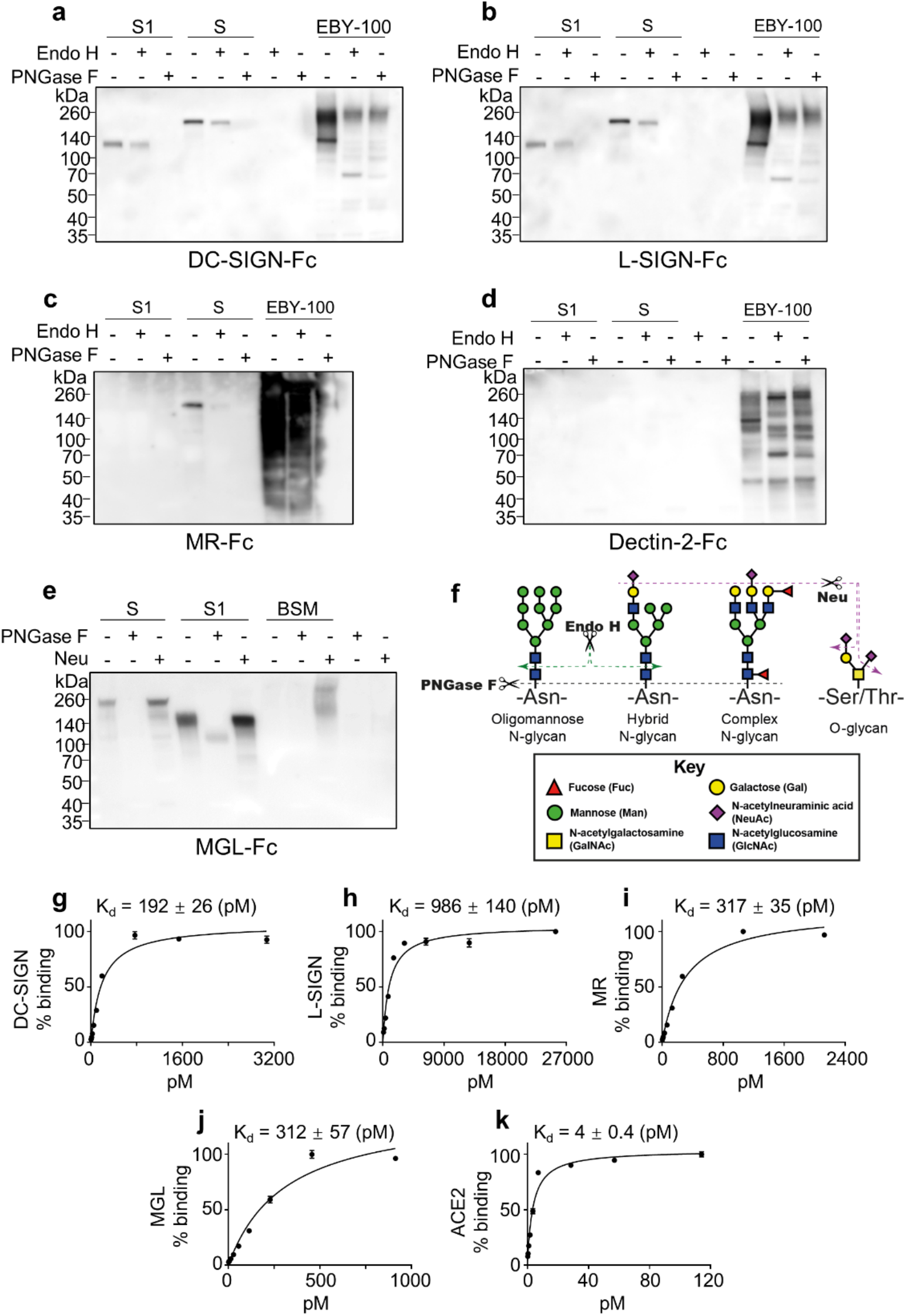
Binding of CLRs to the recombinant SARS-CoV-2 S. (**a-e**) Immunoblots with human Fc-fused CLRs DC-SIGN (**a**), L-SIGN (**b**), MR (**c**), Dectin-2 (**d**) and MGL (**e**) to detect recombinant S1 and S after mock enzymatic digestion or with Endo H, PNGase F or Neu digestion. As negative controls, these glycosidases were also included in some assays. EBY-100 represents the lysates of yeast strain EBY-100. BSM is the recombinant bovine submaxillary mucin. In all assays 5 mM Ca^2+^ was included in solutions of CLRs. (**f**) Schematic presentation of the cleavage sites of Endo H, PNGase F and Neu on N- and O-glycans. Endo H cleaves the oligomannose and hybrid N-glycans, while PNGase F removes all N-glycans including the complex type. Neu removes all sialic acids on N- or O-glycans. (**g-k**) Affinity constant measurement for DC-SIGN (**g**), L-SIGN (**h**), MR (**i**), MGL (**j**) and ACE2 (**k**) by ELISA assay. The plates were coated by recombinant SARS-CoV-2 S trimer. Error bars represent SD of two replicates. The data were plotted as % binding relative to the saturated binding as 100%. In all assays 5 mM Ca^2+^ and Tween-20 were included in solutions of CLRs.

Taken together, our results demonstrate that DC-SIGN and L-SIGN bind the recombinant S via high-mannose and complex N-glycans, while MR recognizes S via its high-mannose moieties only. MGL binding can be largely attributed to N-glycans, but O-glycans could also be partly involved in its recognition.

### CLRs interact with the SARS-CoV-2 S with high affinity

Earlier study showed DC-SIGN and MGL at 1 μg/ml can bind the recombinant receptor binding domain (RBD) of S by ELISA assay^23^. Here, using a native-like trimeric SARS-CoV-2 S protein, we sought to measure the binding affinities of DC-SIGN, L-SIGN, MR and MGL, and the canonical SARS-CoV-2 entry receptor ACE2. The results indicate that the CLRs and ACE2 all bind S in a dose-dependent manner (**Fig. 1g-k**). While ACE2 showed the highest affinity with Kd = 4 pM, binding of DC-SIGN, MGL and MR was strong, with Kd = 192, 312 and 317 pM, respectively. L-SIGN showed moderate affinity with Kd = 986 pM.

### SARS-CoV-2 S glycoforms mediate interactions with CLRs

To directly characterize the nature of S glycans that interact with these CLRs, we performed in-depth N-glycan and O-glycopeptide analyses (**Fig. 2**). A similar set of N-glycans, including oligomannose- and complex-type, was detected in the recombinant full-length S, the S1 and S2, but the relative abundance of each individual glycan varies between samples (**Supplementary Fig. 3**, **Supplementary Table 1**). Oligomannose N-glycans were mainly detected in the full-length S, with the major component being Man5GlcNAc2, serving potential ligand for DC-SIGN, L-SIGN, and MR^24, 25^. Major epitopes on the complex N-glycans were revealed by MALDI-TOF-MS/MS analysis (**Supplementary Fig. 4**). Most complex N-glycans contained core Fuc, and a large proportion were neutral and terminated with either GlcNAc or LacNAc (Gal-GlcNAc) (**Fig. 2**). Notably, our N-glycan microarray analysis revealed that although certain GlcNAc-terminating N-glycans can be bound by DC-SIGN and L-SIGN, the binding was greatly attenuated by the presence of core Fuc (#5 and #7 vs #1 and #3, **Supplementary Fig. 5** and **Supplementary Table 2**). Thus, they are unlikely to be the binding partners of DC-SIGN and L-SIGN. Moreover, MS/MS confirmed the presence of Lewis A/X and LacdiNAc epitopes (GalNAc-GlcNAc) (**Supplementary Fig. 4a & b**). The former was relatively high in the full-length S and the S1, potentially serving as ligands of DC-SIGN and L-SIGN^26, 27^. The latter was particularly abundant in S1 and S2, potentially supporting the binding of MGL^28^. The higher molecular-weight region is dominated by multi-antennary N-glycans with various degrees of sialylation (**Supplementary Fig. 3**), which are not known ligands for those CLRs.

**Fig. 2.**
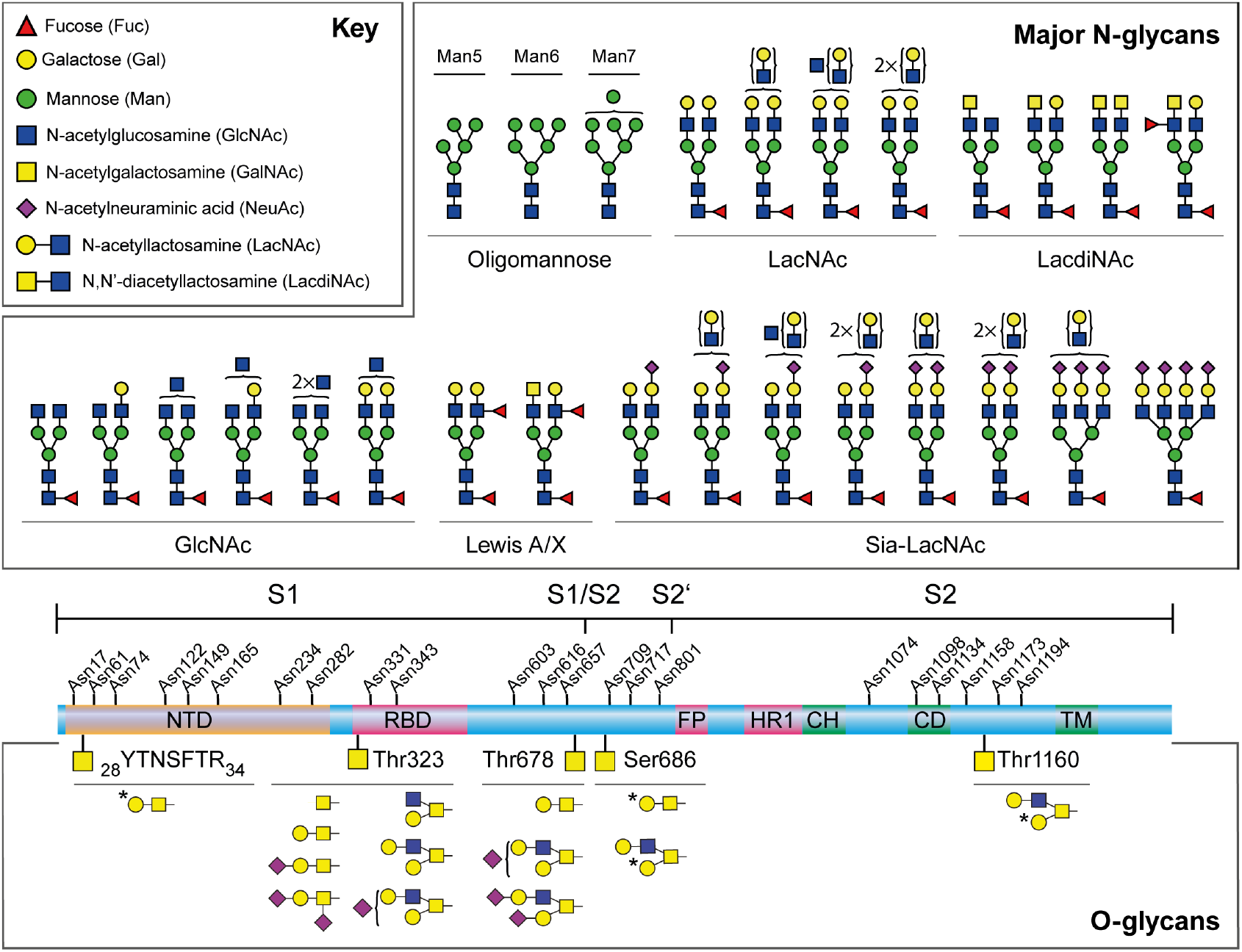
Major N-glycans and O-glycopeptides identified in the recombinant full-length SARS-CoV-2 S. Schematic representation of SARS-CoV-2 S shown in the middle. The positions of N-glycosylation sites are shown on top. Protein domains in the illustration are: N-terminal domain (NTD), receptor-binding domain (RBD), fusion peptide (FP), heptad repeat 1 (HR1), central helix (CH), connector domain (CD), and transmembrane domain (TM). The cleavage sites of S1/S2 and S2’ are labelled. Major N-glycan structures detected by mass spectrometry were categorized by their epitopes on the non-reducing terminal and shown on top. Cartoon symbols above a curly parenthesis indicates sequences corresponding to these compositions cannot be unequivocally defined. The structures presented are only the major glycans on the recombinant full-length S, S1, and S2. A full list of glycans can be found in **Supplementary Table 1**. The O-glycopeptides detected in the full-length S protein are presented at the bottom. The identified O-glycosylation sites are marked on the protein and the O-glycans on each specific site are listed below each site. The LC-MS/MS spectrum of each O-glycopeptide can be found in **Supplementary Fig. 6**. Labelled with asterisk were those found only after neuraminidase treatment.

In an O-glycopeptide-centered analysis of the full-length SARS-CoV-2 S, we identified four new O-glycosylation sites (Tyr28-Arg34, Thr678, Ser 686 and Thr1160) in addition to the previous reported site (Thr323) in the RBD^4, 5^ (**Fig. 2**, **Supplementary Fig. 6** and **Supplementary Table 3**). All sites except Thr323 were partially occupied. Major glycan epitopes identified on these sites include non-, mono- and disialylated core 1 (Galβ1-3GalNAc-R) and core 2 [GlcNAcβ1-6(Galβ1-3)GalNAc-R]. Importantly, Tn antigen (GalNAc), which is a binding determinant for MGL, was only partially present on Thr323. This is in line with our conclusion that MGL mainly binds N-glycans.

Among the four newly discovered O-glycosylation sites, two reside in the furin cleavage site between S1/S2 (Thr678 and Ser686) which are unique to SARS-CoV-2. Although there has not yet been evidence that O-glycosylation plays a role in protease cleavage of S^1, 29^, further investigation is warranted as O-glycosylation does affect protease susceptibility in other systems, as well as antibody recognition^30, 31^.

In summary, our glycomics, glycoproteomics analyses, and glycan microarray analysis confirmed that the glycans on SARS-CoV-2 S could serve as ligands for DC-SIGN, L-SIGN, MR and MGL.

### DC-SIGN and L-SIGN on the cell surface bind SARS-CoV-2 S resulting in its internalization

In order to investigate whether S interacts with the CLRs expressed on cell surfaces, we performed flow cytometry using transduced fibroblast-derived 3T3 cells expressing DC-SIGN and L-SIGN, although the latter had only a L-SIGN+ subpopulation (**Supplementary Fig. 7**). The parental 3T3 cells lacked expression of DC-SIGN or L-SIGN. In accordance with the western blot results, the full-length SARS-CoV-2 S trimer was strongly bound by DC-SIGN+ cells (**Fig. 3a & b**) and to a lesser extent by L-SIGN+ cells (**Fig. 3c & d**). Ten minutes after incubation, DC-SIGN+ cells presented signs of S internalization which was significantly higher than the parental cells after 30 mins (**Fig. 3e**). Thus, our results indicate that SARS-CoV-2 S can be recognized and captured by cells expressing these CLRs, which can lead to internalization of the virus.

**Fig. 3.**
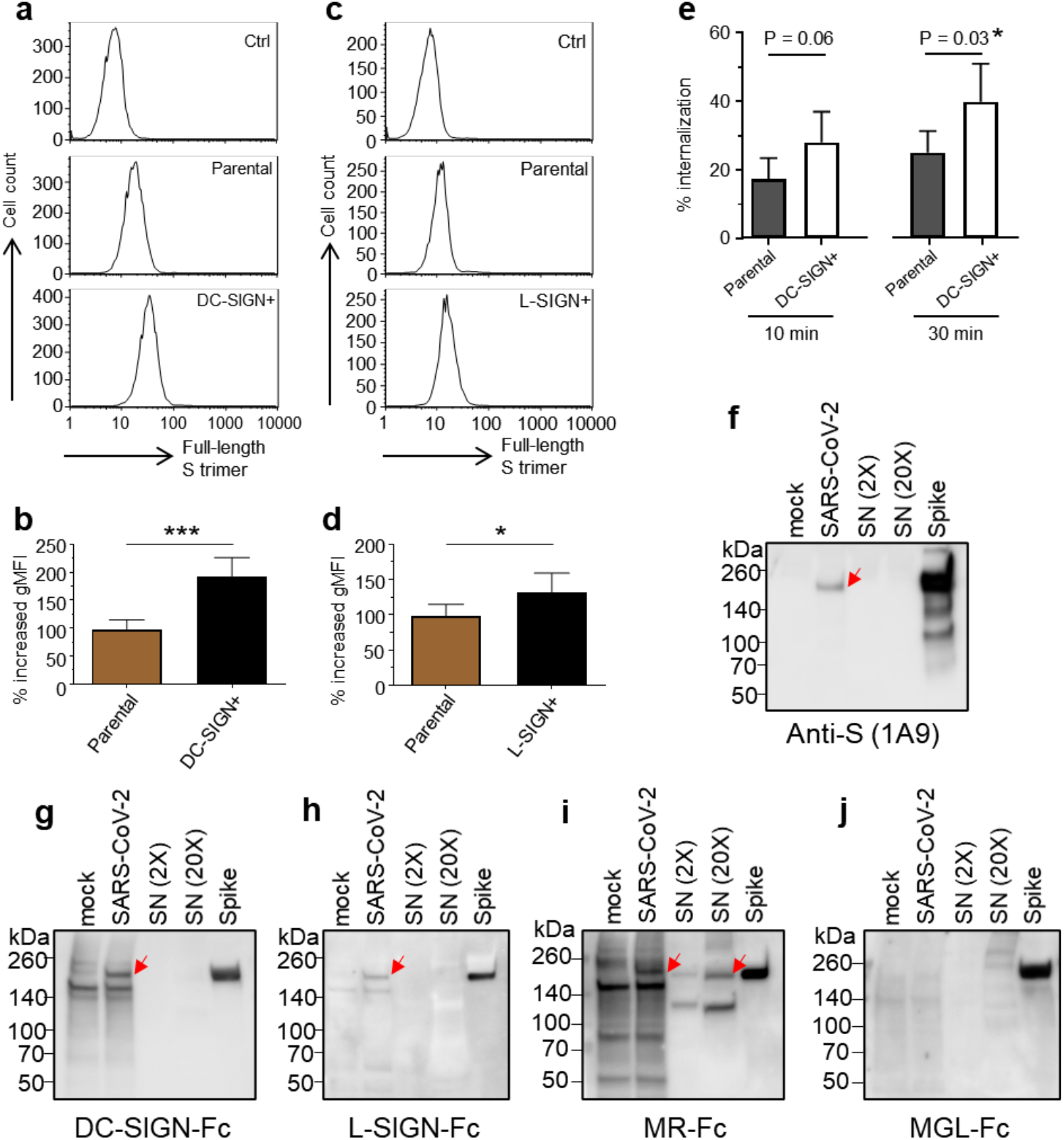
Binding analysis of cells expressing DC-SIGN and L-SIGN and Vero E6 cells infected by SARS-CoV-2. (**a** and **c**) Flow cytometry profiles showing the binding of the full-length SARS-CoV-2 S trimer to parental 3T3 cells (middle), and 3T3-DC-SIGN+ and 3T3-L-SIGN+ cells (bottom). (**b** and **d**) Increased binding of SARS-CoV-2 S trimer to the parental 3T3 cells, the 3T3-DC-SIGN+ and 3T3-L-SIGN+ cells relative to the secondary antibody control. Data presented here is the percentage of increase in geometric mean fluorescence intensity (gMFI). (**e**) Internalization of SARS-CoV-2 S trimer in 3T3-DC-SIGN+ cells compared to the parental 3T3 cells at 10 and 30 min. (**f**-**j**) Immunoblots with anti-S mAb 1A9 (**f**), human Fc-fused CLRs DC-SIGN (**g**), L-SIGN (**h**), MR (**i**), and MGL(**j**) using lysates of Vero E6 cells mock infected (lane 1) or SARS-CoV-2 infected (lane 2), and the SARS-CoV-2-containing culture supernatant (2X and 20X in lane 3 and 4, respectively). Red arrow indicates the bands corresponding to the SARS-CoV-2 S. In all assays 5 mM Ca^2+^ was included in solutions of CLRs. The recombinant SARS-CoV-2 S was used as a positive control.

### CLRs bind S produced by SARS-CoV-2-infected Vero E6 cells

To further explore interactions of a native viral S protein with CLRs, we performed western blot using lysates of SARS-CoV-2-infected Vero E6 cells (**Fig. 3f-j**). Twenty-four hours post infection, SARS-CoV-2 S can be robustly detected in the cell lysates by monoclonal antibody 1A9 (**Fig. 3f**), which binds the recombinant full-length S but not S1 of SARS-CoV-2 (**Supplementary Fig. 7e**). No band was detected in the virus-containing culture supernatant (SN), even at a higher loading amounts (20×).

Among the CLRs, DC-SIGN and L-SIGN recognized a band with identical mobility to the full-length S in the lysates of SARS-CoV-2-infected cells (red arrow, **Fig. 3g & h**). This was not present in the mock-infected cells. MR also exhibited positive binding in the SARS-CoV-2-infected cells, and in addition, in the SN (red arrows, **Fig 3i**). By contrast, MGL, although strongly bound to the recombinant S produced in HEK293 cells, did not appear to bind to S in the Vero E6 cell lysates or in the SN (**Fig. 3j**). These results indicate that the glycosylation status of S is influenced by the source of glycosylation, either in infected cells or the recombinant expression system.

Taken together, our results demonstrate that SARS-CoV-2-infected Vero E6 cells produce S protein which can be bound by DC-SIGN, L-SIGN and MR, suggesting that the virus could be captured by host cells expressing these CLRs via their unique carbohydrate recognition.

### CLRs are expressed on innate immune cells in tissues susceptible to SARS-CoV-2 infection

To characterize the distribution of DC-SIGN, L-SIGN, MR and MGL, we surveyed their expression patterns together with ACE2 in multiple human tissues using public available single-cell RNA sequencing (scRNA-seq) data (**Fig. 4, Supplementary Fig. 8 & 9**). Consistent with previous results^32^, ACE2 expression was generally low in all analyzed datasets, including human lung and upper airway, thymus, pancreas, spleen, ileum, liver and colon (**Fig. 4a & b** and **Supplementary Fig. 8**). In comparison, expression levels of MR and MGL were higher, particularly in cells of the lung and upper airway (**Fig. 4a**). Their expression was mainly restricted to resident immune cells, such as macrophages and dendritic cells, which was also the major cell types producing DC-SIGN, although to a lower extent. L-SIGN was mainly expressed in endothelial cells in liver and pancreas, in lymphatic tissues in ileum, as well as in immune cells, but at much lower levels (**Fig. 4a & b** and **Supplementary Fig. 8**).

**Fig. 4.**
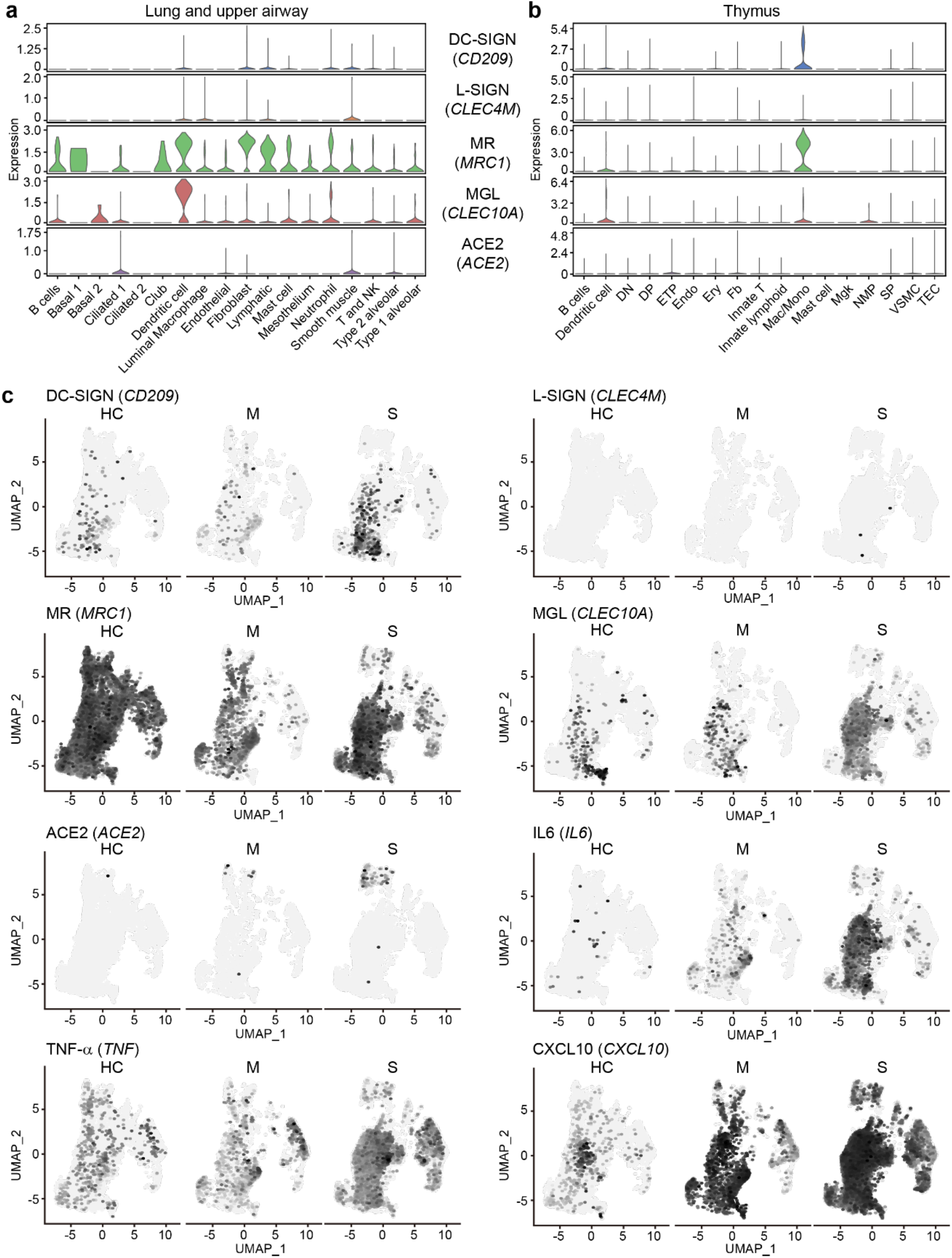
Expression of CLRs in human tissues and CLRs/cytokines/chemokines in BALF from COVID-19 patients. (**a** and **b**) Single-cell transcriptomic analysis of CLRs gene expression (DC-SIGN, L-SIGN, MR, MGL) and ACE2 in lung and upper airway and thymus as indicated. NK, natural killer cells; DN, double-negative T cells; DP, double-positive T cells; ETP, early thymic progenitor; Endo, endothelial cells; Ery, erythrocytes; Fb, fibroblasts; Mono, monocyte; Mac, macrophage; Mgk, megakaryocyte; NMP, neutrophil-myeloid progenitor; SP, single-positive T cells; VSMC, vascular smooth muscle cells; TEC, thymic epithelial cells). (**c**) UMAP showing the gene expression levels of CLRs gene expression (DC-SIGN, L-SIGN, MR, MGL) and ACE2, and selected cytokines and chemokines in BALF immune cells from health controls (HC, n = 3), moderate cases (M, n = 3) and severe/critical cases (S, n = 6).

Emerging evidence showed a strong pathological inflammation in patients with COVID-19, correlated with the presence of monocytes, macrophages and dendritic cells^33, 34^. Therefore, we examined the expression of DC-SIGN, L-SIGN, MR and MGL in available dataset of BALF cells of healthy controls (HC), and patients with moderate (M) and severe (S) COVID-19 (3, 6 and 3 cases, respectively, **Fig. 4c** and **Supplementary Fig. 9**). Consistent with the original report^12^, unbiased clustering identified over 30 distinct cell clusters, with the majority of cells being macrophages (**Supplementary Fig. 9a**). Expression of ACE2 was mainly restricted to epithelial cells of severe patients (**Fig. 4c**). In comparison, MR was predominantly expressed in almost all macrophages and dendritic cells from the three donor groups. Of note, the expression levels of DC-SIGN and MGL were increased in severe COVID-19 patients with elevated amount of proinflammatory monocyte-derived macrophages and inflammatory cytokines and chemokines, including IL-6, TNF, CXCL10, CXCL8, IL-1B, CCL2 and CCL3 (**Fig. 4c** and **Supplementary Fig. 9**). The resulting increased systemic cytokine production from activated macrophages may contribute to the pathophysiology of severe COVID-19.

Our analyses confirm the expression of CLRs and absence of ACE2 in bronchoalveolar and other innate immune cells, indicating that DC-SIGN, MR and MGL might serve as alternative receptors and entry routes in these cells for SARS-CoV-2.

## Discussion

Our results demonstrate that the SARS-CoV-2 S can engage in a glycan-dependent manner with multiple CLRs including DC-SIGN, L-SIGN, MR, and MGL, which are highly expressed in innate immune cells and lymphoid organs, as confirmed by scRNA-seq analysis. The observed interactions with picomolar affinities can initiate receptor-dependent internalization of S as exemplified by DC-SIGN, and potentially provide routes for virus to enter macrophages and dendritic cells. Our scRNA-seq analyses confirmed the co-expression of the CLRs such as DC-SIGN, MR and MGL, and inflammatory cytokines and chemokines in hyperactive macrophages and dendritic cells in patients with COVID-19. This is consistent with the altered cytokine production in SARS-CoV-infected macrophages and dendritic cells, albeit lack productive viral replication^14, 15^. Our results offer a possible explanation to how SARS-CoV-2 spreads to extrapulmonary tissues within the host^35, 36, 37^, as the innate immune cells lacking ACE2 expression can still interalize the virus via those CLRs.

Glycosylation of SARS-CoV-2 S is obviously complex and dependent on the nature of the protein and the glycosylation machinery of infected host cells. Our results indicate that glycosylation potentially determines the immune receptors with which SARS-CoV-2 interact. This variation may lead to virus clearance or on the contrary, results in spread of the virus to other organs, or even other hosts. Interactions of SARS-CoV-2 with CLRs and subsequent internalization suggests a possibility that resident innate immune cells in the lung may lessen the available titer of free virus by endocytosis, thus slowing the onset of symptoms and contributing to asymptomatic or presymptomatic patients^38, 39^. Notably, DC-SIGN and L-SIGN can promote “trans” viral transmission, particularly exemplified in HIV-1^40^. Early studies confirmed that DC-SIGN and L-SIGN-expressing cells were able to transfer SARS-CoV to susceptible target cells^17, 18^. Further research is warranted to explore whether this could occur with SARS-CoV-2. A recent proteomics study discovered strong association between the level of DC-SIGN and variants in the *ABO* locus, the glycosyltransferases required for blood group synthesis, a known genetic risk factor for respiratory failure in COVID-19 ^41^. Given that SARS-CoV-2 RNAemia is detected, particularly in patients with severe disease^38, 42^, it is tempting to speculate that subsequent to CLR-dependent internalization, SARS-CoV-2 could be conveyed by CLR-expressing innate immune cells, and redistributed to permissive tissues where more damage could occur^43^. In accord with this hypothesis, a recent study revealed that SARS-CoV-2 nucleocapsid can be detected in spleens and lymph nodes^44^.

The results here suggest several diagnostic and therapeutic directions. Inhibition of the CLRs that bind virus or inhibition of the glycan-CLR interactions by small molecules might lessen the distribution of virus in COVID-19 patients and potentially limit immune cell (hyper)activation. Analysis of patients for polymorphisms in the genes encoding CLRs might reveal associations with disease severity, as have been found for L-SIGN in SARS-CoV^19^. Finally, defining the differences in glycosylation of S and the variant glycoforms might provide insights into disease severity and spread of the virus in COVID-19 patients.

## Methods

### Recombinant proteins

Recombinant S1 (His-tag), S2 (His-tag), full-length S protein (with mutations R683A and R685A, His-tag), S protein trimer (with mutations R683A and R685A, His-tag), human ACE2 (Fc-tag) were all purchased from Acrobiosystems (S1N-C52H3, S2N-C52H5, SPN-C52H4, SPN-C52H8, AC2-H5257, respectively). The trimeric conformation of the fulllength S protein was validated by the vender using size exclusion chromatography under detection with multi angle light scattering. Human DC-SIGN (Fc-tag), DC-SIGNR (Fc-tag), MGL (Fc-tag) were purchased from Sino Biological (10200-H01H, 10559-H01H, 10821-H01H, respectively). Anti-Spike protein antibody (1A9), which was a mouse monoclonal antibody (IgG1) detecting the spike proteins of both SARS-CoV and SARS-CoV-2 through S2 subunit, was purchased from GeneTex (GTX632604).

Human IgG Fc-tagged MR (MR-Fc), containing the murine C-type lectin domains 4-7 fused to a human IgG Fc-portion, was produced in house. HEK293T cells were transfected with MR-Fc DNA (kind gift from L. Martinez Pomares). Transfection was performed with Lipofectamine LTX with Plus Reagent (Thermo Fisher Scientific) conform with the manufacturer’s guidelines. The cells were incubated with Lipofectamine complex at 37°C in a CO2 incubator for 24 h, and the medium refreshed after 24 h. After 9 days the supernatant containing the produced MR-Fc was collected and purified. MR-Fc was purified from cell culture supernatant using Protein A-agarose beads (Roche). For ELISA experiments, the MR-Fc was purified over mannan-agarose beads (Sigma) and protein A-agarose beads. MR-Fc was quantified by Nanodrop (Thermo Fisher Scientific) and stored at −20°C until further use.

### Cell culture

Human fibroblast cell line 3T3, and the DC-SIGN-, and L-SIGN-transduced 3T3 cells (3T3-DC-SIGN+ and 3T3-DC-SIGNR+, respectively) were obtained through the AIDS Reagent Program, Division of AIDS, NIAID, NIH from Drs. Thomas D. Martin and Vineet N. KewalRamani, HIV Drug Resistance Program, NCI^45^. They were cultured in Dulbecco’s Modified Eagle’s Medium (DMEM) supplemented with 10% (vol/vol) fetal bovine serum (FBS), 1% penicillin-streptomycin and 2 mM glutamine at 37°C in 5% CO2. Vero E6 cells (ATCC CRL-1586) were cultured in DMEM supplemented with 2 mM L-glutamine, 50 units/ml penicillin, 50 mg/ml streptomycin, and 10% FBS. Infection studies were performed in cell culture medium supplemented with 2% FBS.

The Saccharomyces strain EBY-100 was purchased from ATCC and cultured under the recommended condition by the vendor. The EBY-100 cell extracts were prepared with the lysis buffer (20 mM Tris, 100 mM NaCl, 1 mM EDTA, 2% Triton X-100, 1% SDS, pH 7.6). The protein concentration was quantified by Pierce™ BCA Protein Assay Kit (Thermo Fisher Scientific).

### Preparation of SARS-CoV-2

All work with infectious SARS-CoV-2, isolate USA_WA1/2020 was performed at the National Emerging Infectious Diseases Laboratories, Boston University under Biosafety Level 4. Vero E6 cells seeded in 6-well plates were infected with SARS-CoV-2 at a multiplicity of infection (MOI) of 1 or mock infected. After 1 h of virus adsorption at 37°C in 5% CO2, the inocula were removed and replaced with DMEM supplemented with 2% FBS. Twenty-four hours post-infection, cell supernatants were removed, the cells were scraped and pelleted by low-speed centrifugation. Cell pellets were washed once with PBS and resuspended in 270 μl Cell Extraction Buffer (Thermo Fisher Scientific). A 900 μl aliquot of a SARS-CoV-2 stock (Titer: 1.58 × 10^7^ TCID50 units/ml) was also used for analysis. Both the cell lysates and the viral stock were inactivated by addition of SDS (1 % final concentration) and boiling for 10 minutes prior to removal from the Biosafety Level 4 facility. The inactivated cell lysates and virus stock were aliquoted and stored at −80°C until further use. The protein concentration was quantified by Pierce™ BCA Protein Assay Kit (Thermo Fisher Scientific).

### Western Blot and Lectin Blot

The recombinant S, S1 and S2 proteins, together with controls were subject to enzymatic digestion by PNGase F (P0708L, New England Biolabs), Endo H (P0702L, New England Biolabs), neuraminidase (10269611001, Roche) or mock treatment before loading onto the SDS-PAGE gel. The enzymatic digestion was following the protocols recommended by the venders. Recombinant S, S1 and S2 protein and BSM were used at 1μg/lane, while the EBY-100 extract was used at 15 μg/lane. The reaction mixture was directly loaded on the 4-20% SDS-PAGE gel (GenScript) following addition of 4× laemmli loading buffer. The cell extracts either from SARS-CoV-2 infected or mock infected Vero E6 cells (both at 50 μg protein/lane), together with the culture supernatant (at 12 μg (2×) and 120 μg (20×) protein/lane), were loaded on the SDS-PAGE gel following addition of 4× laemmli loading buffer and 10 min heating. The gels were either stained with Coomassie Brilliant Blue or Colloidal Blue (both from Thermo Fisher Scientific) to visualize the proteins, or transferred to a PVDF membrane (Thermo Fisher Scientific) for Western blots.

The Western blot and lectin blot analysis were performed following the protocols published previously^46^. The proteins were under the following concentrations: DC-SIGN, L-SIGN, Dectin-2, MGL all at 1 μg/ml, MR at 2.5 μg/ml, biotinylated plant lectins GNA and VVA both at 1 μg/ml, Con A at 0.1 μg/ml, or antibodies mAb100 at 1 μg/ml and anti-spike antibody 1A9 at 0.5 μg/ml. The CLRs were prepared either in BSA-TTBS buffer [1% BSA in 20 mM Tris, 300 mM NaCl, 5 mM CaCl2, 2 mM MgCl2, pH 7.4, with 0.05% Tween-20] or BSA-TTBS-EDTA [1% BSA in 20 mM Tris, 300 mM NaCl, 10 mM EDTA, pH 7.4) with 0.05% Tween-20 (TTBS)] to check the calcium dependence. HRP-labelled secondary reagents including goat anti-human IgG-HRP, goat anti-mouse IgG-HRP (Jackson ImmunoResearch Laboratories) or streptavidin-HRP (Vector Laboratories) are all used in 1:10,000 dilution.

### ELISA

The ELISA assay was performed following the protocols published previously^46^. The S-protein trimer was immobilized in a 96-well plate (0.2 μg/well) overnight at 4°C. The proteins were serial diluted (1:2) and applied to each well. HRP-labeled goat anti-human IgG was used at 1:1,000 dilution and the color development was with TMB ELISA Substrate (Abcam, ab171523) The experiment was performed in duplicate and the background subtraction was done by subtracting the corresponding binding signals obtained with BSA-immobilized wells. Affinity constant was calculated with GraphPad Prism 6.0 (GraphPad Software, Inc.).

### Flow cytometry

For cell surface binding assay, the cultured 3T3, 3T3-DC-SIGN and 3T3-L-SIGN cells were collected and washed with cold PBS once. The cells were incubated with full-length S protein trimer (20 μg/ml) or the monoclonal antibody against DC-SIGN, #120507 (2 μg/ml) and L-SIGN, #120604 (2 μg/ml), or buffer only (negative control) on ice for 30 minutes. Binding of the full-length S protein was detected by mouse anti-His IgG (Invitrogen MA121315, 2 μg/ml) and Alexa Fluor-488-labelled goat anti-mouse IgG (Invitrogen A11001, 1:500).

Binding of anti-DC-SIGN and anti-L-SIGN were detected directly with Alexa Fluor 488-labelled goat anti-mouse IgG. The results shown were from three independent experiments analyzed by FlowJo software.

### Internalization assay

To measure internalization, the cells were suspended and incubated with S protein trimer on ice as stated above. The cells were then centrifuged and fixed in 0.5% paraformaldehyde (T0) in binding buffer or allowed for further incubation at 37 °C for 10 or 30 min (Tn) and ended with 0.5% paraformaldehyde. Fixed cells were stained with mouse anti-His IgG and Alexa Fluor-488-labelled goat anti-mouse IgG as indicated above for residual S trimer detection. Internalization rate (%) was calculated by the formula: [gMFI of S trimer(T0)-gMFI of S trimer (Tn)]/gMFI of S trimer (T0) × 100. The results shown were from five independent experiments with comparable results.

### Microarray analysis

Microarray analyses on the complex N-glycan array were conducted as reported previously^47^. The microarray slides were probed with Fc-tagged DC-SIGN and L-SIGN at 10 μg/ml diluted in 1% BSA (TSM with 0.05% Tween) and binding was detected with Alexa Fluor 488 labelled goat anti-human IgG (H+L) (Invitrogen) at 5 μg/ml.

### Glycomics analysis

The full length S, S1 and S2 proteins were subjected to N-glycomics analysis following the protocols published previously^48^ with some modifications. Briefly, 20 μg each of the recombinant proteins were lyophilized and digested with PNGase F (P0701S, New England Biolabs Inc.) according to the manufacture’s instruction. Post digestion, the glycans were purified by Sep-Pak C18 cartridges (WAT054955, Waters Corp.) and subjected to permethylation. The permethylated glycans were purified by chloroform extraction and Sep-Pak C18 cartridges prior to mass spectrometric analysis.

MALDI-TOF MS and MALDI-TOF/TOF MS/MS analysis of the permethylated glycans were performed with the UltrafleXreme mass spectrometer (Bruker Corp.) equipped with a Smartbeam II laser. Spectrum between mass-to-charge (m/z) 1000 and 5500 was acquired under reflectron positive mode. Selective peaks were subject to MS/MS analysis. Each MS spectrum presented an accumulation of 20,000 laser shots.

### O-glycan LC-MS analysis

Three microgram of full-length S protein were treated with PNGase F (NEB) for N-glycan release according to the manufacture’s instruction, followed by optional sialic acid removal using 0.02 U sialidase (Roche), prior to in-gel trypsin treatment^49^. After overnight trypsin treatment an optional AspN (Roche) digestion was performed in 1:25 ratio (enzyme:protein). The samples were dried down in a speed vac concentrator, reconstituted in Cal and further diluted in 0.1 % FA for subsequent LC-MS analysis. All samples were prepared in triplicates.

LC-MS was performed using an Ultimate 3000 nano LC coupled to an Orbitrap Fusion Lumos mass spectrometer (both from Thermo Fisher). Three microliters of each sample were loaded onto a C18 precolumn (C18 PepMap 100, 300 μm x 5 mm, 5 μm, 100 Å, Thermo Fisher Scientific) with 15 μL/min solvent A (0.1% FA in H2O) for 3 min and separated on a C18 analytical column (picofrit 75 μm ID x 150 mm, 3 μm, New Objective) using a linear gradient of 2 % to 45 % solvent B (80% acetonitrile, 0.1% FA) over 106 min at 400 nL/min. The ion source was set at 2100 V spray voltage and 200 °C ion transfer tube temperature. MS scans were performed in the orbitrap at a resolution of 60000 within a scan range of m/z 600 - m/z 1600, a RF lens of 30%, AGC target of 1e5 for a maximum injection time of 50 ms. The top 15 precursors were selected for MS2 in a data dependent manner, within a mass range of *m/z* 600 – 1600 and a minimum intensity threshold of 5e4 and an isolation width of 1.5 *m/z*. HCD was performed at 27 % collision energy and detected in the orbitrap with a resolution of 30000 with the first mass at *m/z* 120, an AGC target of 2e5 and a maximum injection of 250 ms.

EThcD spectra were acquired in a product ion-dependent manner ([HexNAc+H]^+^-ion) based on the method above. Precursor isolation width was set to 1.2 *m/z*. Calibrated charge-dependent ETD parameters were used with supplemental activation collision energy of 25%, an AGC target of 2e5 and a maximum injection time of 350 ms.

Glycopeptide identification was performed using Byonic version 3.5 (Protein Metrics Inc.). Trypsin and/or AspN were set as proteases with a maximum of two missed cleavage sites. Mass tolerance for the precursor and HCD fragments was 10 ppm and for EThcD 20 ppm. The glycan database was “O-glycan 78 mammalian”. The following modifications were allowed: carbamidomethyl (Cys; fixed), oxidation (Met; variable common 2), pyroglutamine on N-term (Gln, variable, rare 1), acetylation N-term (variable rare1), deamidation (Asn, variable common 2), formylation N-term (variable rare1). Glycopeptides with a score above 250 were selected and further manually inspected.

### scRNA-sequencing

For the expression of ACE2 and receptor genes across different tissues, datasets were retrieved from published datasets in multiple human tissues, including lung and upper airway^50, 51^, ileum^52^, colon^53^, pancreas^54^, spleen^55^ and thymus^56^. These datasets are available and can be visualized and assessed through a website portal (www.covid19cellatlas.org)^32^. The processed .h5ad files were loaded by “read_h5ad” and violin plots were illustrated by “pl.stacked_violin” in scanpy 1.5.1, which is a model for single cell analysis in Python^57^. For the single-cell RNA sequencing analysis of bronchoalveolar immune cells in patients with SARS-CoV-2, dataset was retrieved from Liao et al.^12^ and Gene Expression Omnibus (GEO) under the accession number GSE145926, which contains 6 severe and 3 moderate SARS-CoV-2 patients and 3 healthy controls^12^.

The raw data with .h5 format was loaded for analysis through R package Seurat v3^58, 59^. For each sample, cells were filtered out if they contain genes less than 200 or more than 6000, or if their UMI is less than 1000, or mitochondrial gene percentage larger than 0.1. The remaining cells were integrated into a gene-barcode matrix and then normalized and log-transformed. We identified 2000 highly variable genes by ‘vst’ method for the downstream PCA analysis. RNA count and the percent of mitochondrial genes were regressed out in the scaling step. We chose the top 50 principal components for the UMAP and graph-based clustering. The cell type identity was referred to Liao et al.^12^ and feature genes were demonstrated by “DimPlot”.

### Data availability

All data is available in the manuscript or in the supplementary information. The single cell RNA-sequencing datasets are available online with the corresponding references.

## Acknowledgments

This work was supported by National Institutes of Health grants P41GM103694 and R24 GM137763, and Evergrande MassCPR subaward 280870.5116795.0025 to E.M., as well as an NIH supported (1UL1TR001430) COVID-19 Related Research Award from the Boston University Clinical and Translational Science Institute (CTSI) to A.E.. SARS-CoV-2 isolate USA_WA1/2020 was kindly provided by CDC’s Principal Investigator Natalie Thornburg, and the World Reference Center for Emerging Viruses and Arboviruses (WRCEVA). The following reagent was obtained through the NIH AIDS Reagent Program, Division of AIDS, NIAID, NIH: NIH 3T3, NIH 3T3-DC-SIGN+ and NIH 3T3-L-SIGN+ from Drs. Thomas D. Martin and Vineet N. KewalRamani. We thank Dr. Jamie Heimburg-Molinaro for help in editing the manuscript.

## Author contributions

C.G. and R.D.C. conceived the project. C.G. and Y.M. performed the western blots and ELISA. N.J. and K.S. performed N-glycomics and O-glycoproteomics analysis. J.Z. maintained the cell culture and performed cell binding assays and internalization assay. C.G. performed glycan microarray analysis. H.Z. and J.L. performed scRNA-seq analysis. A.J.H., E.M., I.D. and J.K. provided important materials for the experiments. K.T., A.E. and R.D.C. supervised experiments. C.G. wrote the first draft of the manuscript and all authors contributed to the final version.

## Additional information

Supplementary Information is available for this paper.

## Supplementary Information

Supplementary Figures 1-9

Supplementary Tables 1-3 (in separate excel files)

## References

1. Hoffmann M, et al. SARS-CoV-2 Cell Entry Depends on ACE2 and TMPRSS2 and Is Blocked by a Clinically Proven Protease Inhibitor. Cell 181, 271–280 e278 (2020).

2. Monteil V, et al. Inhibition of SARS-CoV-2 Infections in Engineered Human Tissues Using Clinical-Grade Soluble Human ACE2. Cell 181, 905–913.e907 (2020).

3. Letko M, Marzi A, Munster V. Functional assessment of cell entry and receptor usage for SARS-CoV-2 and other lineage B betacoronaviruses. Nat Microbiol 5, 562–569 (2020).

4. Shajahan A, Supekar NT, Gleinich AS, Azadi P. Deducing the N- and O-glycosylation profile of the spike protein of novel coronavirus SARS-CoV-2. Glycobiology, 2020.2004.2001.020966 (2020).

5. Watanabe Y, Allen JD, Wrapp D, McLellan JS, Crispin M. Site-specific glycan analysis of the SARS-CoV-2 spike. Science, 2020.2003.2026.010322 (2020).

6. Drouin M, Saenz J, Chiffoleau E. C-Type Lectin-Like Receptors: Head or Tail in Cell Death Immunity. Front Immunol 11, 251 (2020).

7. van Kooyk Y. C-type lectins on dendritic cells: key modulators for the induction of immune responses. Biochem Soc Trans 36, 1478–1481 (2008).

8. van Kooyk Y, Rabinovich GA. Protein-glycan interactions in the control of innate and adaptive immune responses. Nat Immunol 9, 593–601 (2008).

9. Routhu NK, et al. Glycosylation of Zika Virus is Important in Host-Virus Interaction and Pathogenic Potential. International journal of molecular sciences 20, 5206 (2019).

10. Gringhuis SI, den Dunnen J, Litjens M, van der Vlist M, Geijtenbeek TBH. Carbohydrate-specific signaling through the DC-SIGN signalosome tailors immunity to Mycobacterium tuberculosis, HIV-1 and Helicobacter pylori. Nature Immunology 10, 1081–1088 (2009).

11. Guan W-j, et al. Clinical Characteristics of Coronavirus Disease 2019 in China. New England Journal of Medicine 382, 1708–1720 (2020).

12. Liao M, et al. Single-cell landscape of bronchoalveolar immune cells in patients with COVID-19. Nature Medicine, (2020).

13. Tian S, Hu W, Niu L, Liu H, Xu H, Xiao S-Y. Pulmonary Pathology of Early-Phase 2019 Novel Coronavirus (COVID-19) Pneumonia in Two Patients With Lung Cancer. Journal of Thoracic Oncology 15, 700–704 (2020).

14. Law HKW, et al. Chemokine up-regulation in SARS-coronavirus–infected, monocyte-derived human dendritic cells. Blood 106, 2366–2374 (2005).

15. Tseng C-TK, Perrone LA, Zhu H, Makino S, Peters CJ. Severe Acute Respiratory Syndrome and the Innate Immune Responses: Modulation of Effector Cell Function without Productive Infection. The Journal of Immunology 174, 7977–7985 (2005).

16. Jeffers SA, et al. CD209L (L-SIGN) is a receptor for severe acute respiratory syndrome coronavirus. Proceedings of the National Academy of Sciences of the United States of America 101, 15748–15753 (2004).

17. Marzi A, et al. DC-SIGN and DC-SIGNR Interact with the Glycoprotein of Marburg Virus and the S Protein of Severe Acute Respiratory Syndrome Coronavirus. Journal of Virology 78, 12090–12095 (2004).

18. Yang Z-Y, et al. pH-Dependent Entry of Severe Acute Respiratory Syndrome Coronavirus Is Mediated by the Spike Glycoprotein and Enhanced by Dendritic Cell Transfer through DC-SIGN. Journal of Virology 78, 5642–5650 (2004).

19. Chan VSF, et al. Homozygous L-SIGN (CLEC4M) plays a protective role in SARS coronavirus infection. Nature Genetics 38, 38–46 (2006).

20. Shih Y-P, et al. Identifying Epitopes Responsible for Neutralizing Antibody and DC-SIGN Binding on the Spike Glycoprotein of the Severe Acute Respiratory Syndrome Coronavirus. Journal of Virology 80, 10315–10324 (2006).

21. Han DP, Lohani M, Cho MW. Specific Asparagine-Linked Glycosylation Sites Are Critical for DC-SIGN- and L-SIGN-Mediated Severe Acute Respiratory Syndrome Coronavirus Entry. Journal of Virology 81, 12029–12039 (2007).

22. Goh JB, Ng SK. Impact of host cell line choice on glycan profile. Crit Rev Biotechnol 38, 851–867 (2018).

23. Chiodo F, et al. Novel ACE2-Independent Carbohydrate-Binding of SARS-CoV-2 Spike Protein to Host Lectins and Lung Microbiota. bioRxiv, 2020.2005.2013.092478 (2020).

24. Guo Y, et al. Structural basis for distinct ligand-binding and targeting properties of the receptors DC-SIGN and DC-SIGNR. Nat Struct Mol Biol 11, 591–598 (2004).

25. Martinez-Pomares L. The mannose receptor. Journal of Leukocyte Biology 92, 1177–1186 (2012).

26. Feinberg H, Mitchell DA, Drickamer K, Weis WI. Structural Basis for Selective Recognition of Oligosaccharides by DC-SIGN and DC-SIGNR. Science 294, 2163–2166 (2001).

27. Noll AJ, et al. Human DC-SIGN Binds Specific Human Milk Glycans. Biochemical Journal, (2016).

28. Mortezai N, et al. Tumor-associated Neu5Ac-Tn and Neu5Gc-Tn antigens bind to C-type lectin CLEC10A (CD301, MGL). Glycobiology 23, 844–852 (2013).

29. Walls AC, Park Y-J, Tortorici MA, Wall A, McGuire AT, Veesler D. Structure, Function, and Antigenicity of the SARS-CoV-2 Spike Glycoprotein. Cell 181, 281–292.e286 (2020).

30. de las Rivas M, et al. Molecular basis for fibroblast growth factor 23 O-glycosylation by GalNAc-T3. Nature Chemical Biology 16, 351–360 (2020).

31. Silver ZA, et al. Discovery of O-Linked Carbohydrate on HIV-1 Envelope and Its Role in Shielding against One Category of Broadly Neutralizing Antibodies. Cell Reports 30, 1862–1869.e1864 (2020).

32. Sungnak W, et al. SARS-CoV-2 entry factors are highly expressed in nasal epithelial cells together with innate immune genes. Nature Medicine 26, 681–687 (2020).

33. Merad M, Martin JC. Pathological inflammation in patients with COVID-19: a key role for monocytes and macrophages. Nature Reviews Immunology 20, 355–362 (2020).

34. Vabret N, et al. Immunology of COVID-19: current state of the science. Immunity, (2020).

35. Wang W, et al. Detection of SARS-CoV-2 in Different Types of Clinical Specimens. JAMA 323, 1843–1844 (2020).

36. Wu Y, et al. Prolonged presence of SARS-CoV-2 viral RNA in faecal samples. The Lancet Gastroenterology & Hepatology 5, 434–435 (2020).

37. Lamers MM, et al. SARS-CoV-2 productively infects human gut enterocytes. Science, eabc1669 (2020).

38. Huang C, et al. Clinical features of patients infected with 2019 novel coronavirus in Wuhan, China. The Lancet 395, 497–506 (2020).

39. Arons MM, et al. Presymptomatic SARS-CoV-2 Infections and Transmission in a Skilled Nursing Facility. New England Journal of Medicine 382, 2081–2090 (2020).

40. Geijtenbeek TBH, et al. DC-SIGN, a Dendritic Cell–Specific HIV-1-Binding Protein that Enhances trans-Infection of T Cells. Cell 100, 587–597 (2000).

41. Brufsky A, Lotze MT. DC/L-SIGNs of Hope in the COVID-19 Pandemic. Journal of Medical Virology n/a.

42. Hogan CA, et al. High frequency of SARS-CoV-2 RNAemia and association with severe disease. medRxiv, 2020.2004.2026.20080101 (2020).

43. Li H, et al. SARS-CoV-2 and viral sepsis: observations and hypotheses. The Lancet 395, 1517–1520 (2020).

44. chen y, et al. The Novel Severe Acute Respiratory Syndrome Coronavirus 2 (SARS-CoV-2) Directly Decimates Human Spleens and Lymph Nodes. medRxiv, 2020.2003.2027.20045427 (2020).

45. Wu L, Martin TD, Vazeux R, Unutmaz D, KewalRamani VN. Functional Evaluation of DC-SIGN Monoclonal Antibodies Reveals DC-SIGN Interactions with ICAM-3 Do Not Promote Human Immunodeficiency Virus Type 1 Transmission. Journal of virology 76, 5905–5914 (2002).

46. Matsumoto Y, et al. Identification of Tn antigen O-GalNAc-expressing glycoproteins in human carcinomas using novel anti-Tn recombinant antibodies. Glycobiology 30, 282–300 (2019).

47. Gao C, et al. Unique Binding Specificities of Proteins toward Isomeric Asparagine-Linked Glycans. Cell chemical biology 26, 535–547 e534 (2019).

48. Jia N, et al. The Human Lung Glycome Reveals Novel Glycan Ligands for Influenza A Virus. Scientific Reports 10, 5320 (2020).

49. Plomp R, et al. Site-Specific N-Glycosylation Analysis of Human Immunoglobulin E. Journal of proteome research 13, 536–546 (2014).

50. Deprez M, et al. A single-cell atlas of the human healthy airways. bioRxiv, 2019.2012.2021.884759 (2019).

51. Vieira Braga FA, et al. A cellular census of human lungs identifies novel cell states in health and in asthma. Nature Medicine 25, 1153–1163 (2019).

52. Martin JC, et al. Single-Cell Analysis of Crohn’s Disease Lesions Identifies a Pathogenic Cellular Module Associated with Resistance to Anti-TNF Therapy. Cell 178, 1493–1508.e1420 (2019).

53. Smillie CS, et al. Intra- and Inter-cellular Rewiring of the Human Colon during Ulcerative Colitis. Cell 178, 714–730.e722 (2019).

54. Baron M, et al. A Single-Cell Transcriptomic Map of the Human and Mouse Pancreas Reveals Inter- and Intra-cell Population Structure. Cell Systems 3, 346–360.e344 (2016).

55. Madissoon E, et al. scRNA-seq assessment of the human lung, spleen, and esophagus tissue stability after cold preservation. Genome Biology 21, 1 (2019).

56. Park J-E, et al. A cell atlas of human thymic development defines T cell repertoire formation. Science 367, eaay3224 (2020).

57. Wolf FA, Angerer P, Theis FJ. SCANPY: large-scale single-cell gene expression data analysis. Genome Biol 19, 15 (2018).

58. Stuart T, et al. Comprehensive Integration of Single-Cell Data. Cell 177, 1888–1902 e1821 (2019).

59. Butler A, Hoffman P, Smibert P, Papalexi E, Satija R. Integrating single-cell transcriptomic data across different conditions, technologies, and species. Nat Biotechnol 36, 411–420 (2018).

