## Supplementary Figure 1,2,3,4,5,6,7,8,9 for "SARS-CoV-2 Spike Protein Interacts with Multiple Innate Immune Receptors"


**Supplementary Information:**

Supplementary Figures 1-9

Supplementary Tables 1-3 (in separate excel files)

a

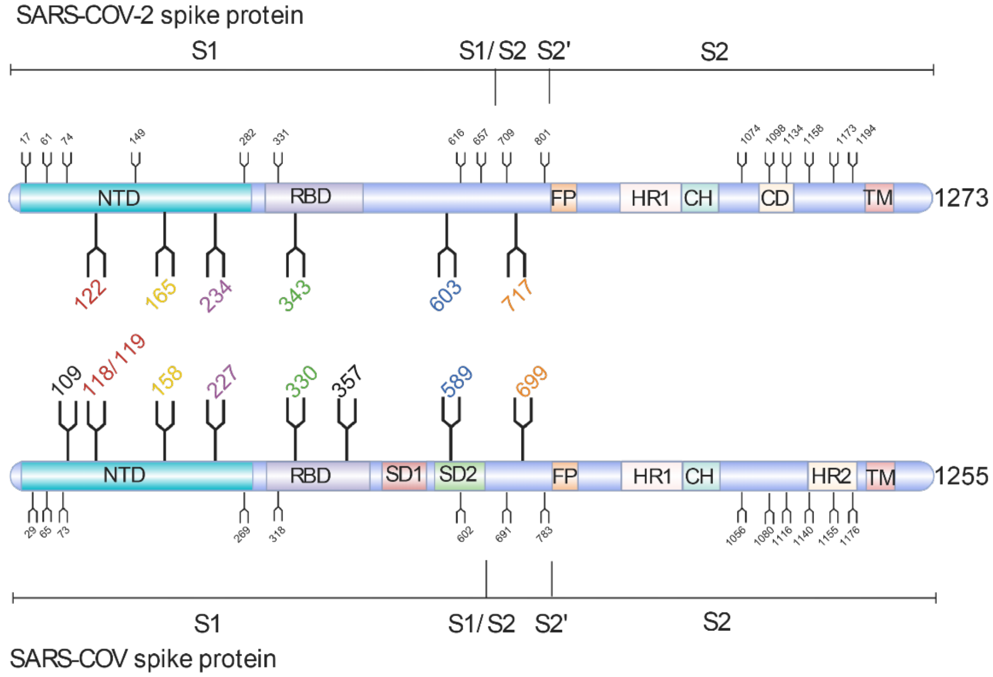


b

| Site of SARS-CoV-2 | 112 | 122 | 165 | 234 | 343 | 370 | 603 | 717 |
|---|---|---|---|---|---|---|---|---|
| Glycan profile | None | M5<br>hybrid<br>FA2/FA1B | FA3/FA2B<br>FA2/FA1B<br>A3/A2B<br>A2/A1B | M8<br>M9<br>M7<br>M6<br>M5 | FA2/FA1B<br>FA3/FA2B<br>FA1<br>hybrid<br>M5 | None | M5<br>FA2/FA1B | M5<br>M6<br>M7<br>hybrid |
| Site of SARS-CoV | 109 | 118/119 | 158 | 227 | 330 | 357 | 589 | 699 |
| Glycan profile | FA3/FA2B<br>FA2/FA1B<br>FA4/FA3B | FA2/FA1B<br>FA3/FA2B<br>FA4/FA3B | M7<br>M6<br>M5<br>M8 | M9<br>M8 | FA2/FA1B<br>FA3/FA2B | FA2/FA1B<br>Fhybrid<br>M5<br>hybrid<br>FA3/FA2B | M5<br>hybrid<br>M6<br>A1 | FA4/FA3B<br>A3/A2B |

**Supplementary Fig. 1 Comparison of the S glycoprotein of SARS-CoV-2 and SARS-CoV**.

**(a)** The N-glycosylation sites are shown on the protein. The cleavage sites of S1/S2 and S2' were also labelled. Eight sites (109, 118/119, 158, 227, 330, 357, 589 and 699) which were previously reported to be involved in DC-SIGN and L-SIGN binding[1,2], were highlighted on the SARS-CoV S. The corresponding conserved sites (122, 165, 234, 343, 603 and 717) in the SARS-CoV-2 S were also highlighted for comparison.

**(b)** N-glycan profiles on each site highlighted on the S glycoproteins of SARS-CoV-2 and SARS-CoV are listed in the table. Data is as shown in refs[3] and[4]. Abbreviations: M is mannose, F is fucose, A is antenna, B is bisecting.

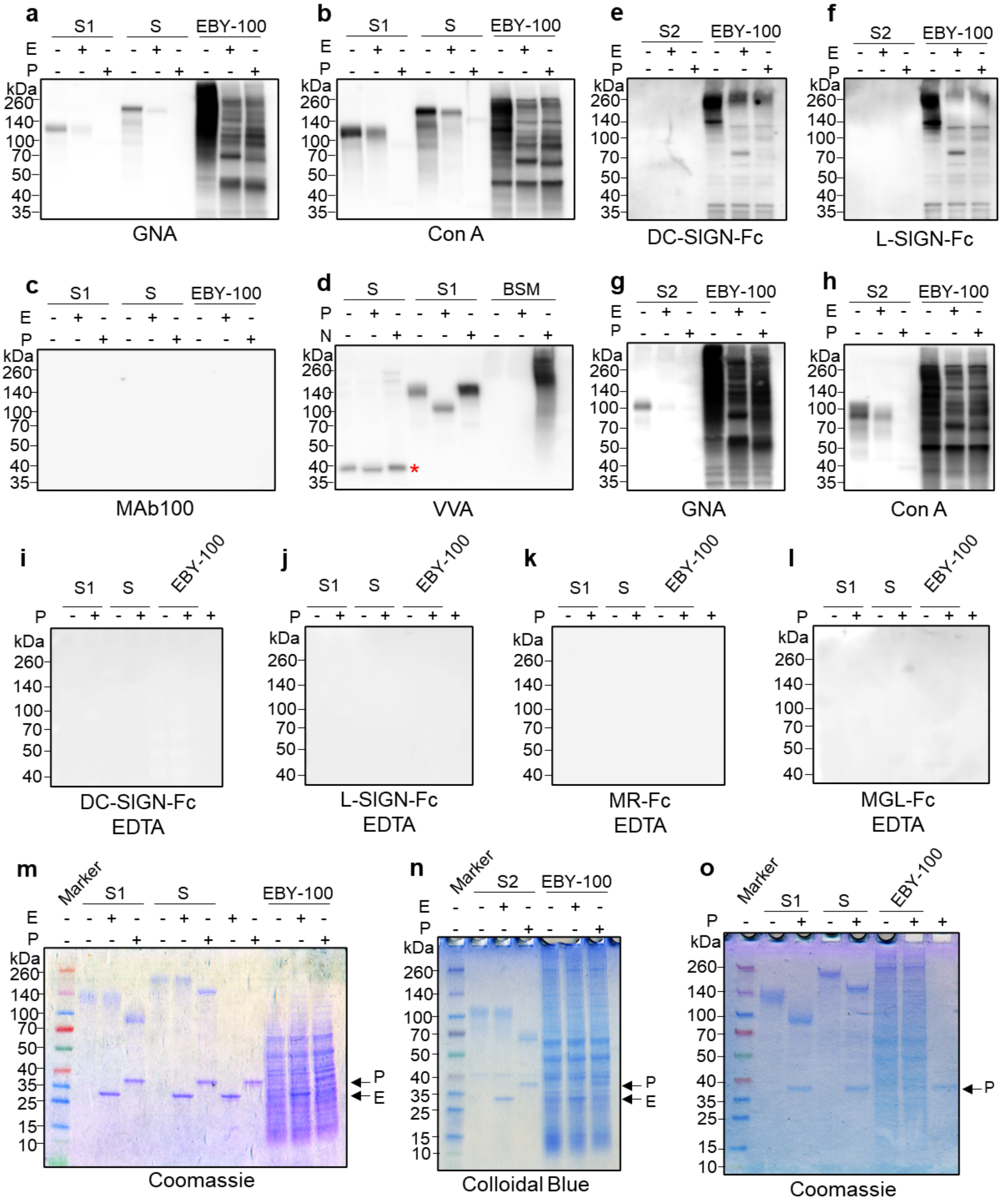


**Supplementary Fig. 2 Binding of CLRs to the recombinant SARS-CoV-2 S, S1 and S2.**

(**a-d**) Immunoblots with biotinylated plant lectins GNA (**a**), ConA (**b**) and VVA (**d**), and a monoclonal antibody mAb100 (**c**) to detect the recombinant SARS-CoV-2 S1 and S with mock enzymatic digestion or with Endo H (E), PNGase F (P) or neuraminidase (N) digestion. Monoclonal antibody mAb100 is known to have a specificity for $Man_3GlcNAc_2$ glycan structures but here showed no binding. Labelled with asterisked in (**d**) are bands not characterized in the SARS-CoV-2 S preparation. This could be due to the fragmentation or unknown impurities. In all assays 5 mM $Ca^{2+}$ was included in solutions of CLRs.

(**e-h**) Immunoblots with human IgG Fc-fused CLRs DC-SIGN (**e**), L-SIGN (**f**), and biotinylated plant lectins GNA (**g**), and Con A (**h**) to detect recombinant SARS-CoV-2 S2 with mock enzymatic digestion or with Endo H (E) or PNGase F (P) digestion. In all assays 5 mM $Ca^{2+}$ was included in solutions of CLRs.

(**i-l**) Immunoblots with human IgG Fc-fused CLRs DC-SIGN (**i**), L-SIGN (**j**), MR (**k**), and MGL (**l**) to recombinant SARS-CoV-2 S1 and S with mock enzymatic digestion or with PNGase F (P) digestion. In all experiments EDTA was included with no $Ca^{2+}$ in the solutions of CLRs.

(**m-o**) Coomassie Brilliant Blue or Colloidal Blue staining of the corresponding SDS-PAGE gels. Arrows indicate the positions of PNGase F (P) and endo H (E).

In the experiments, lysates of EBY-100 or the recombinant bovine submaxillary mucin (BSM) were used as controls.

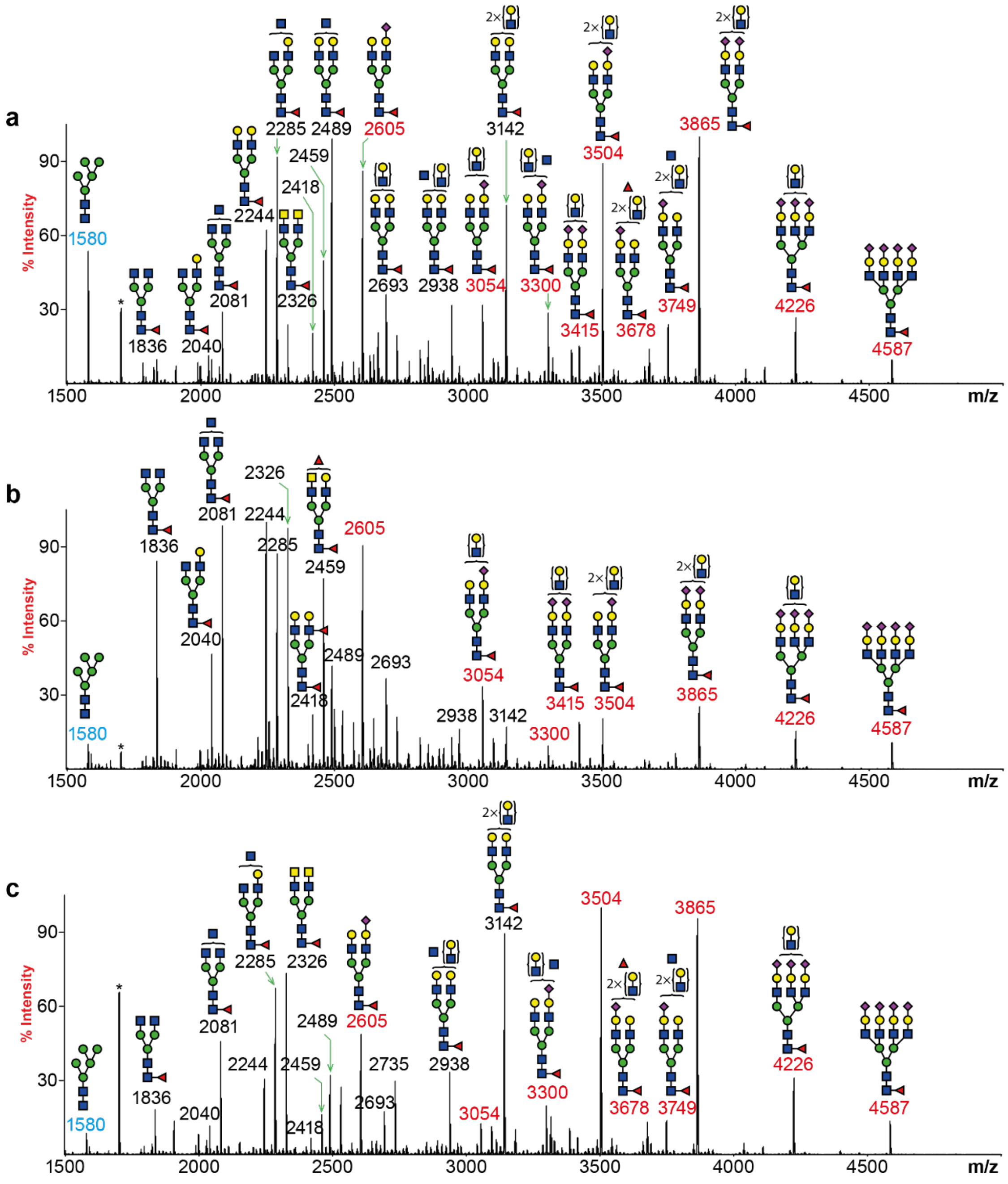


**Supplementary Fig. 3 N-glycan mass spectrometry profiles of the SARS-CoV-2 Spike proteins**.

(**a-c**) MALDI-TOF-MS spectra of the full-length S (**a**), the S1 (**b**) and the S2 (**c**) glycoproteins. All molecular ions detected represent permethylated species and are present in the form of $[M+Na]^+$. Peaks are color-coded based on their structural features in which oligomannose structures are in cyan, non-sialylated glycans are in black and sialylated glycans are in red.

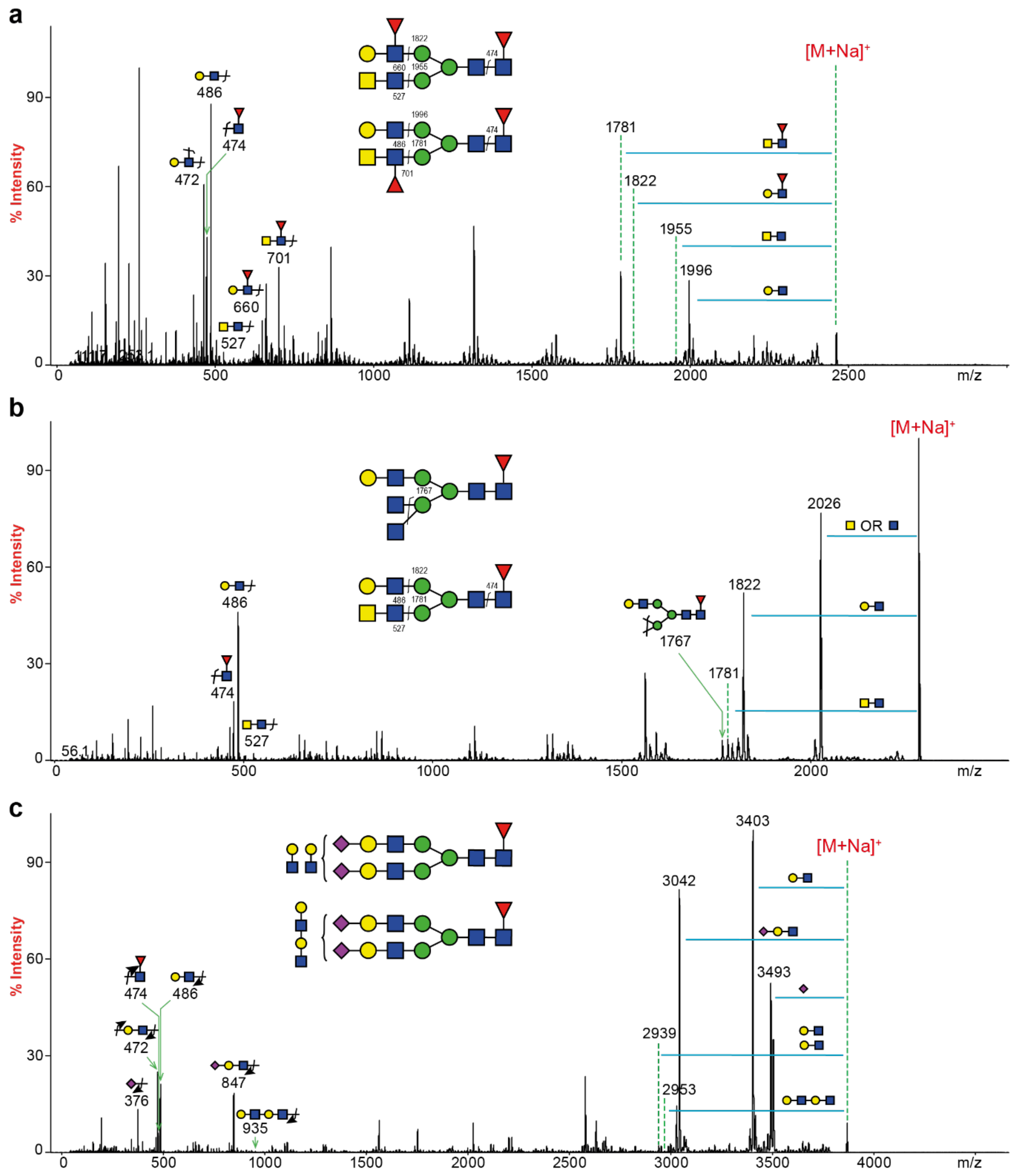


**Supplementary Fig. 4 MS/MS analyses of the SARS-CoV-2 S.** All molecular ions detected represent permethylated species and are present in the form of [M+Na]$^{+}$. In all three MS/MS spectra, the expression of core-fucose was confirmed by the fragment ion at *m/z* 474.

(**a**) MALDI-TOF/TOF-MS/MS analysis of a permethylated N-glycan at *m/z* 2459. Two components with either a fucosylated GlcNAc (LeA/X) or a fucosylated LacdiNAc were confirmed. The fragment ions at *m/z* 660 and 701 indicated the presence of a LeA/X or a fucosylated LacdiNAc epitope, respectively. Their corresponding Y-ions can be detected at *m/z* 1822 and 1781, respectively.

(**b**) MALDI-TOF/TOF-MS/MS analysis of a permethylated N-glycan at *m/z* 2286. Two components with either a terminal GlcNAc or a LacdiNAc were confirmed. The fragment ion at *m/z* 1767 was generated from double-cleavage fragmentation, which was resulted from the losses of two terminal GlcNAc residues. The fragment ion at *m/z* 527 represented a terminal LacdiNAc motif and its corresponding Y-ion was detected at *m/z* 1781.

(**c**) MALDI-TOF/TOF-MS/MS analysis of a permethylated N-glycan at *m/z* 3865. The MS/MS spectrum identified both tetra- and triantennary complex N-glycans. The composition of the fragment ion at *m/z* 2939 represented a YY-ion that was generated by the double cleavage of two terminal LacNAc units. It indicated that the molecular ion was a tetra-antennary N-glycan. A tri-antennary isomeric form was also observed. The detection of the B-ion at *m/z* 935 and its corresponding Y-ion at 2953 suggested the presence of a repeating LacNAc motif with two LacNAc units sequentially connected. The presence of an elongated Sialylated LacNAc motif was not detected.

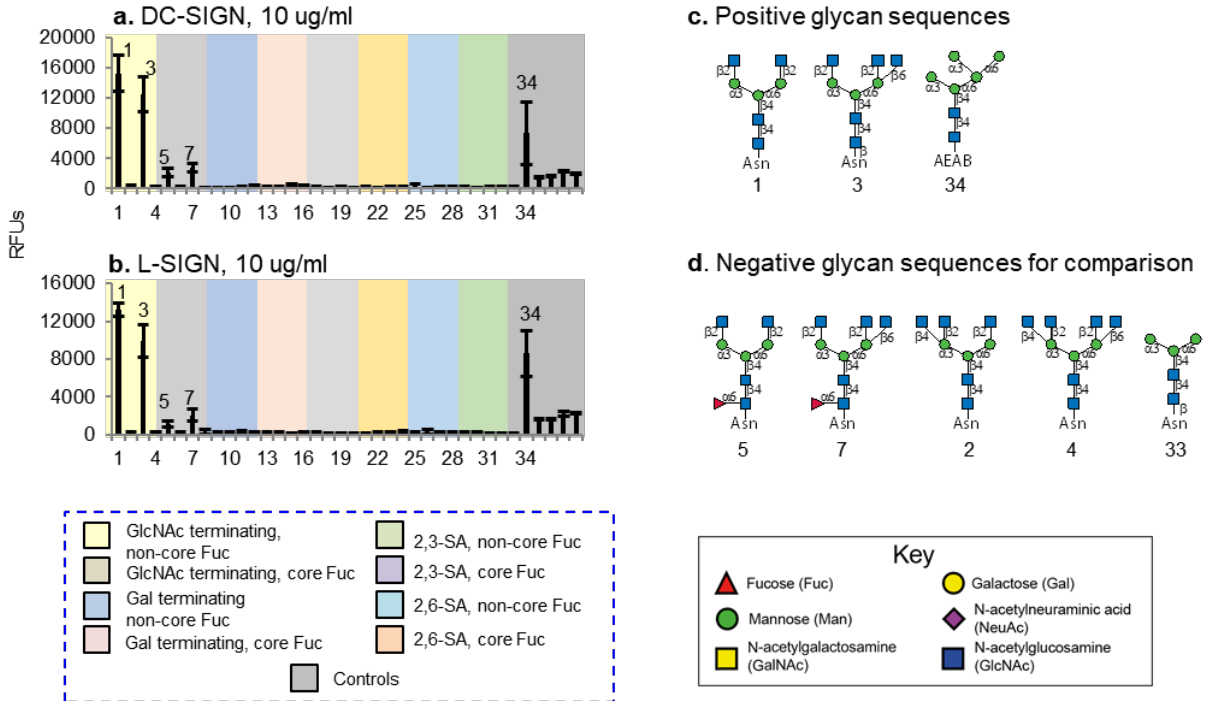


**Supplementary Fig. 5 Microarray analyses of DC-SIGN and L-SIGN using the N-glycan array.**

(**a** and **b**) Microarray binding profile of DC-SIGN (**a**) and L-SIGN (**b**). Results are shown as relative fluorescence units (RFUs) by averaging the background-subtracted fluorescence signals of 4 replicate spots, error bars represent the standard deviation among the 4 values. The RFUs and the standard deviation were listed in Supplemental Table S3. The probes are grouped as indicated in the colored panels.

(**c**) Sequences of glycans that are positively bound on the array. The numbers correspond to the probe numbers in the chart in (**a**) and (**b**).

(**d**) Sequences of selected glycans that are not bound on the array for comparisons. The numbers correspond to the probe numbers in the chart in (**a**) and (**b**).

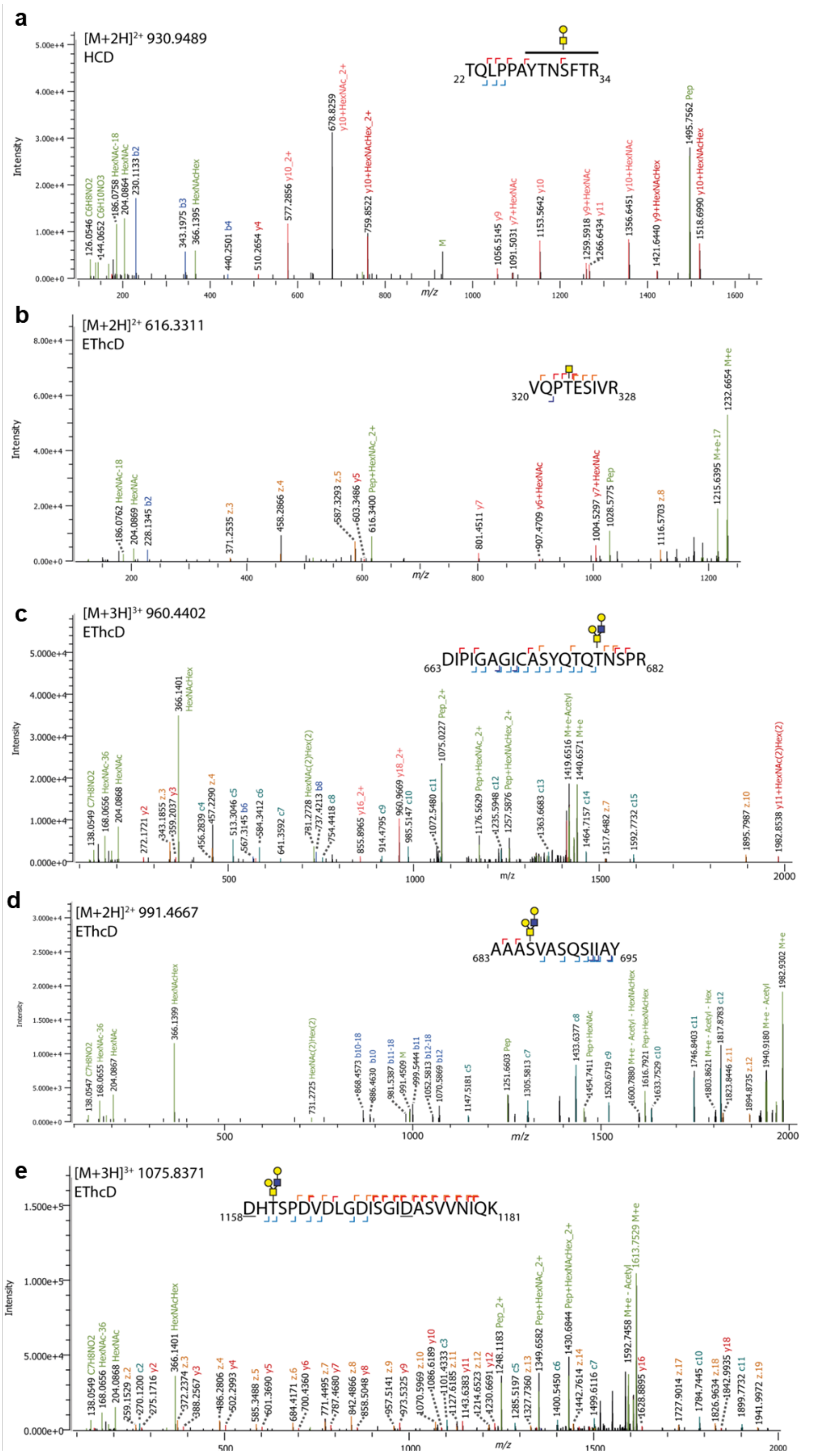


**Supplementary Fig. 6 S protein O-glycosylation sites identified by LC-MS/MS analysis.**

(**a**) O-glycosylation site between Tyr28-Arg34. HCD spectrum of the trypsin and sialidase treated O-glycopeptide at *m/z* 930.9489 $[M+2H]^{2+}$ corresponding to ${}_{22}$TQLPPAYTNSFTR${}_{34}$ with the O-glycan HexNAc$_1$Hex$_1$. Due to the peak y7+HexNAc the O-glycosylation site is assigned between Tyr28-Arg34.

(**b**) O-glycosylation site Thr323. EThcD spectrum of the tryptic O-glycopeptide at *m/z* 616.3311 $[M+2H]^{2+}$ corresponding to ${}_{320}$VQPTESIVR${}_{328}$ with the O-glycan HexNAc$_1$.

(**c**) S protein O-glycosylation site Thr678. EThcD spectrum of the trypsin, AspN and sialidase treated O-glycopeptide at *m/z* 960.4402 $[M+3H]^{3+}$ corresponding to ${}_{63}$DIPIGAGICASYQTQTNSPR${}_{682}$ containing the O-glycan HexNAc$_2$Hex$_2$.

(**d**) O-glycosylation site Ser686. EThcD spectrum of the trypsin and sialidase treated O-glycopeptide at *m/z* 991.4667 $[M+2H]^{2+}$ corresponding to ${}_{683}$AAASVASQSIIAY${}_{695}$ containing the O-glycan HexNAc$_2$Hex$_2$. The recombinant protein contains two single mutations in this region, R683A and R685A, which potentially could alter the O-glycosylation of the natural form.

(**e**) O-glycosylation site Thr1160. EThcD spectrum of the trypsin and sialidase treated O-glycopeptide at *m/z* 1075.8371 $[M+3H]^{3+}$ corresponding to ${}_{1158}$$\underline{\text{D}}$HTSPDVDLGDISGI$\underline{\text{D}}$ASVVNIQK${}_{1181}$ ($\underline{\text{D}}$ indicated deamidation after PNGase F N-glycan release) containing the O-glycan HexNAc$_2$Hex$_2$.

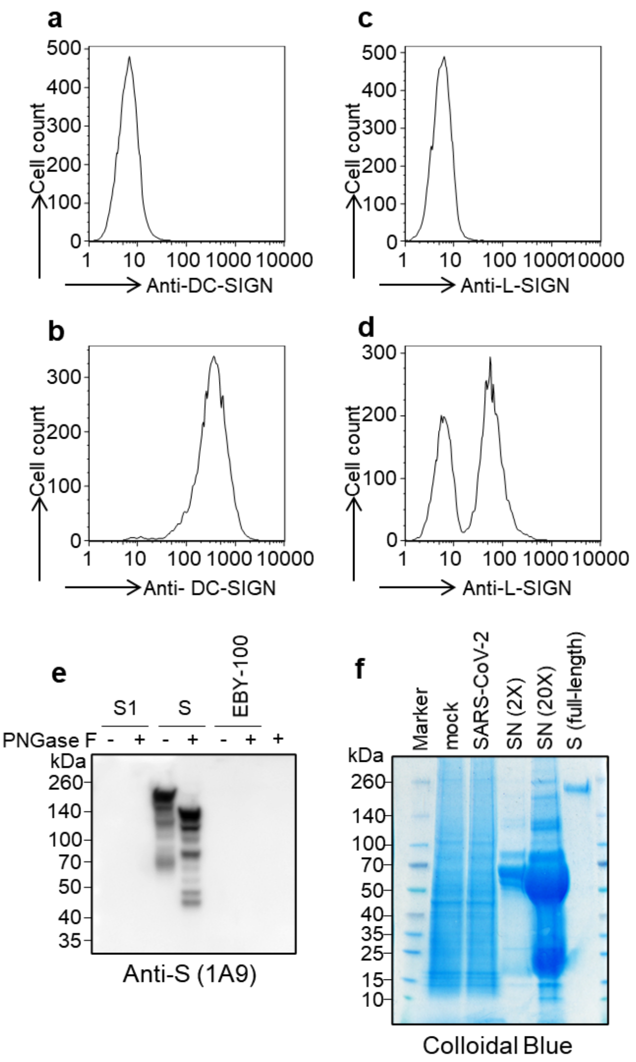


**Supplementary Fig. 7 Flow cytometry analysis of cells expressing DC-SIGN and L-SIGN and immunoblots with recombinant S1 and S and Vero E6 cells infected by SARS-CoV-2**.

(**a** and **b**) FACS analysis with anti-DC-SIGN using 3T3 cells (**a**) and 3T3-DC-SIGN+ cells (**b**).

(**c** and **d**) FACS analysis with anti-L-SIGN using 3T3 cells (**c**) and 3T3-DC-SIGN+ cells (**d**).

(**e**) Western blot with anti-S mAb 1A9 to detect recombinant S1 and S with mock enzymatic digestion or with PNGase F digestion. EBY-100 represents the lysates of yeast strain EBY-100.

(**f**) Colloidal Blue staining of the SDS-PAGE gel using lysates of Vero E6 cells mock infected (lane 2) or SARS-CoV-2 infected (lane 3), and the SARS-CoV-2 virus-containing culture supernatant (2X and 20X in lane 4 and 5, respectively).

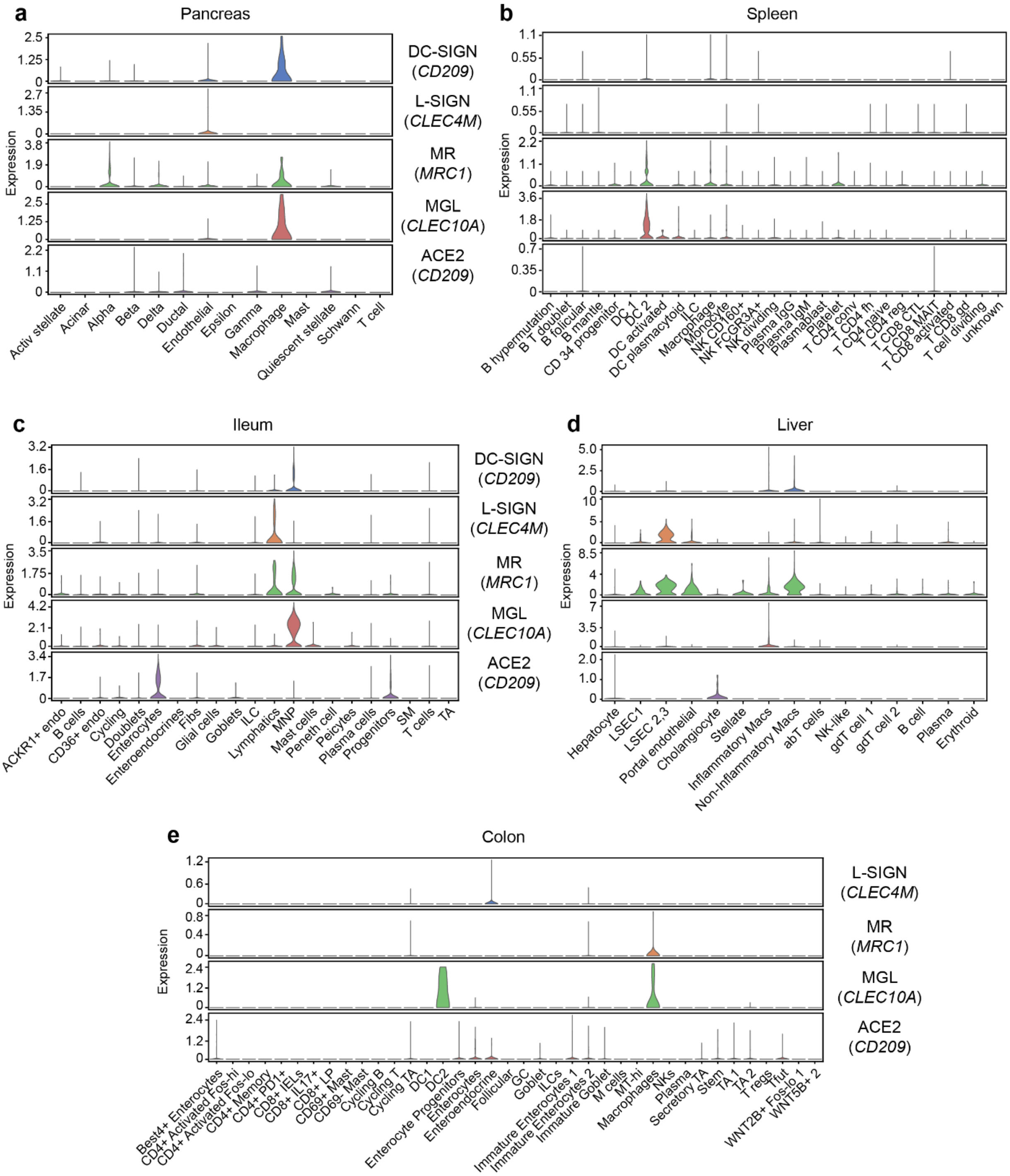


**Supplementary Fig. 8 Expression of CLRs and ACE2 in various human tissues**.

(**a** to **e**) Single-cell transcriptomic analysis of CLRs gene expression (DC-SIGN, L-SIGN, MR, MGL) and ACE2 in pancreas, spleen, ileum, liver and colon as indicated. NK, natural killer cells; DC, dendritic cells; Endo, endothelial cells; ILC, innate lymphoid cells; MNP, mononuclear phagocytes; Fibs, fibroblasts; SM, smooth muscle; TA, transit-amplifying cells; LSEC, liver sinusoidal endothelial cells; Macs, macrophages.

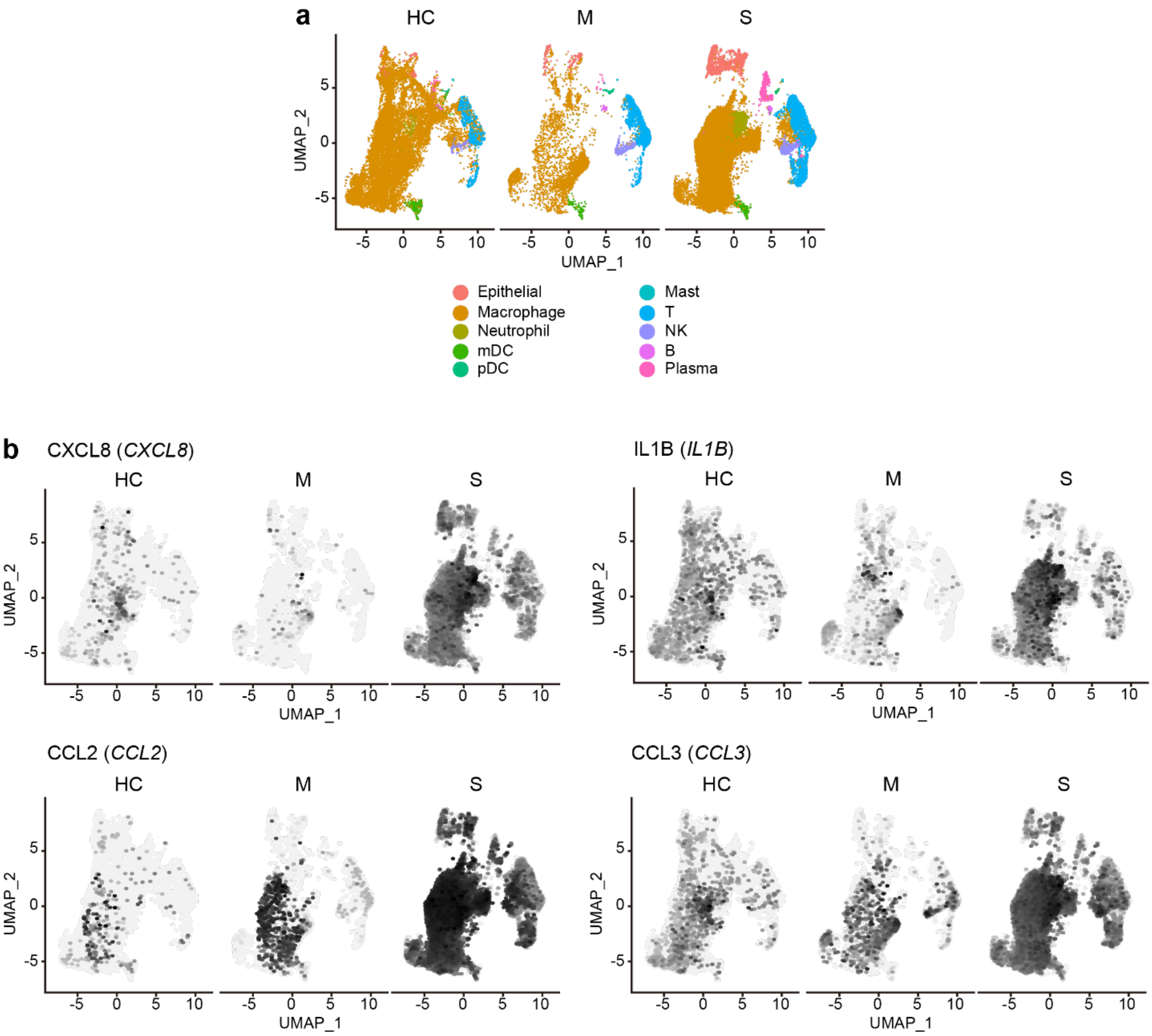


**Supplementary Fig. 9 Single-cell transcriptomic analysis of bronchoalveolar immune cells in patients with COVID-19.**

(**a**) UMAP showing the unbiased clustering of major cell types and associated clusters in BALFs from health controls (HC, n = 3), moderate cases (M, n = 3) and severe/critical cases (S, n = 6). NK, natural killer cells; mDC, myeloid dendritic cells; pDC, plasmacytoid dendritic cells.

(**b**) UMAP showing the gene expression levels of selected cytokines and chemokines including CXCL8, IL1B, CCL2 and CCL3 in BALF.

In separate excel files:

**Supplementary Table 1.** N-glycan composition SARS-CoV-2

**Supplementary Table 2.** Microarray analysis of DC-SIGN and L-SIGN on N-glycan array

**Supplementary Table 3.** O-glycosylation overview

**References:**


1 Shih, Y.-P. *et al.* Identifying Epitopes Responsible for Neutralizing Antibody and DC-SIGN Binding on the Spike Glycoprotein of the Severe Acute Respiratory Syndrome Coronavirus. *Journal of virology* **80**, 10315-10324, doi:10.1128/jvi.01138-06 (2006).

2 Han, D. P., Lohani, M. & Cho, M. W. Specific Asparagine-Linked Glycosylation Sites Are Critical for DC-SIGN- and L-SIGN-Mediated Severe Acute Respiratory Syndrome Coronavirus Entry. *Journal of virology* **81**, 12029-12039, doi:10.1128/jvi.00315-07 (2007).

3 Watanabe, Y. *et al.* Vulnerabilities in coronavirus glycan shields despite extensive glycosylation. *bioRxiv*, 2020.2002.2020.957472, doi:10.1101/2020.02.20.957472 (2020).

4 Watanabe, Y., Allen, J. D., Wrapp, D., McLellan, J. S. & Crispin, M. Site-specific glycan analysis of the SARS-CoV-2 spike. *Science*, 2020.2003.2026.010322, doi:10.1126/science.abb9983 (2020).
